# Learning genetic values of individuals with incomplete pedigree, genomic and phenotypic data

**DOI:** 10.1101/2025.09.11.675626

**Authors:** Daniel Gianola, Ignacio Aguilar, Olga Ravagnolo

## Abstract

Prediction of outcomes is important in personalized medicine, animal and plant breeding. Typical inputs for building prediction models in agriculture include genealogies, molecular markers and phenotypes. Information is seldom complete, i.e., there may be individuals that lack at least one of such inputs. For instance, all individuals may possess pedigree data but only a fraction is genotyped for molecular markers. A solution for such situation is known as “single-step best linear unbiased prediction” (SS-BLUP). A more general scenario is one where, in addition to the setting of SS-BLUP, there are subjects with genomic data but lacking genealogy, with or without phenotypes. Our study presents a novel “single-step” prediction method that accommodates a wider degree of incompleteness than SS-BLUP. It does not employ imputation or approximations and is based on basic Bayesian principles of combining distinct prior opinions. The proposed method, Hy-BLUP (“Hy” for “hybrid”) uses a prior that combines knowledge from the population about variation derived from pedigree and from markers, as if these two sources of information were independent. Such assumption may over-state prior precision, but Bayesian theory dictates that it should be over-ridden as information from data accrues.The Bayesian logic defines the weights assigned to the sources implictly. However, additional weights (*w*_A_and *w*_G_ for pedigree and genomic information, respectively) may be introduced as tuning parameters. From an inferential perspective, the weights and the variance components are not jointly identified in the likelihood function. However, given the variance components, some Bayesian learning about the weights can be obtained. The prior induces a precision matrix (inverse of the covariance matrix) automatically, without use of cumbersome matrix algebra arguments or approximations. The prior is combined with the data and, given the weights (if any) and variance parameters, the estimating “mixed model” equations can be built and computed directly. The method was evaluated using a publicly available data set consisting of 599 inbred lines of wheat genotyped for binary markers and with full pedigree information; the target trait was grain yield. The evaluation used several experiments that simulated various patterns of incompleteness and a training-testing layout supplemented by bootstrapping or random reconstruction of sets. There were minor differences between SS-BLUP and Hy-BLUP in predictive ability.The discussion includes a multiple-trait generalization of Hy-BLUP that may be useful in situations where some individuals are not phenotyped for some trait (e.g., animal carcass weight in a fully-pedigreed breeding nucleus) while others (not pedigreed) are genotyped, scored and destroyed for commercial or laboratory purposes. The study provides a proof-of-concept of the potential usefulness of Hy-BLUP for routine genome-enabled prediction in individuals with irregular patterns of information.

## 1 Introduction

Prediction is important in personalized medicine, plant and animal breeding. Typically, a model is fitted (trained) to data to deliver point and interval predictions conveying an expectation and a measure of uncertainty about the yet-to be-observed outcomes. Predictor variables are, e.g., biomarkers, and outcomes may be some clinical status or a set of quantitative traits in a crop or animal species. Our focus is on quantitative traits.

Henderson (1973, 1975) probably represents the first model-based cohesive theoretical framework for prediction in a quantitative genetics setting. He addressed the problem of learning a vector of unobserved breeding values given genealogical and phenotypic data, and later (Henderson 1977) considered situations where some individuals possess pedigree data but lack records of performance. Henderson examined three settings differing with respect to the degree of knowledge about the joint distribution of observables and unobservables. The best predictor in the smallest mean squared error sense is the conditional (given observables) expectation function (Henderson 1973; Bulmer 1980; Fernando and Gianola 1986). The least restrictive situation discussed by Henderson, i.e., unknown first moments and second moments known, led to the development of best linear unbiased prediction (BLUP). It was found later that BLUP could be interpreted from a Bayesian perspective (Dempfle 1977; Gianola and Fernando 1986), which enabled important extensions of the prediction paradigm to make it suitable for genomic data, (Meuwissen et al. 2001; Gianola et al. 2003; Gianola 2013). The era of “genomic selection” or “genome-enabled prediction” started soon after Meuwissen et al. (2001), with a marked impact on animal and plant breeding and more recently in human medicine (e.g., Yang et al. 2011; de los Campos et al. 2010, 2013).

At the onset of genomic selection most animals and plants had full genealogical and phenotypic information but only a few had molecular data, due to cost. The picture changed later for several species, notably dairy cattle, where millions of animals now possess molecular marker information. In beef cattle or sheep raised under low input or pastoral conditions, however, it is still the case that only a fraction of individuals is genotyped. If all individuals had marker information, pedigree-based BLUP could be replaced by “genomic” BLUP or GBLUP. To accommodate scenarios where genotyping is far from complete, several researchers developed “single-step” or ssGBLUP (e.g., Misztal et al. 2009; Aguilar et al. 2010; Christensen and Lund 2010). Essentially, ssGBLUP imputes unknown breeding values of individuals without markers but with pedigree, from those with markers. ssGBLUP represented a significant breakthrough because all pedigreed animals, with or without genotypes, could be used in genetic evaluation while retaining the logic, flexibility and computational feasibility of BLUP.

A more general scenario, however, is one where, in addition to the setting in ssGBLUP, there are some individuals that possess genomic data and unknown genealogy, with or without phenotypes. The objective of our study is to present a “single-step” prediction method that accommodates a wider degree of incompleteness than ssGBLUP. The proposed method, referred to as Hy-BLUP (“Hy” stands for “hybrid”) follows a Bayesian logic and does not employ an imputation of genomic values from a pedigree, as done in ssGBLUP.

The point of departure in Hy-BLUP is that there are two sources of information (Bayesian “opinions”) about the breeding value of an individual, *a priori*. The first source represents beliefs from an infinitesimal model: prior uncertainty about breeding values is conveyed by 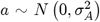 where 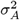 is the additive genetic variance encoded by the infinitesimal model of inheritance and *N* (.) is a Gaussian distribution. The second source posits 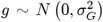, *a priori*; here, 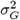 is the additive genomic variance captured by a possibly large but finite number of markers. Each of the two distributions represents a distinct prior opinion about the unknown “true” breeding value, which is defined at the level of unobservable segregating quantitative trait loci or QTL (de los Campos et al. 2015). Variation in genotypes at the QTL loci generates the true unknown additive genetic variance. The infinitesimal and genomic models are distinct representations of the unknown breeding value. Our approach combines the two pieces of prior uncertainty with information conveyed by the available phenotypic records, following ideas of Robertson (1955) and Dempfle (1977) and in a manner consistent with the Bayesian view (Gianola and Fernando 1986; Sorensen and Gianola 2002).

The manuscript is organized as follows. First, an Excursus on ssGBLUP is used to motivate the problem. Second, Hy-BLUP is developed in a step-wise manner with toy examples presented in Supplementary File S1. Third, a publicly available wheat data set is employed to illustrate Hy-BLUP and how it differs from ABLUP (“A” stands for pedigree-based), GBLUP (genomic BLUP) and ssGBLUP. The paper concludes with a discussion.

## 2 Excursus: ssGBLUP

A standard representation of a single-trait mixed linear model under Gaussian assumptions is

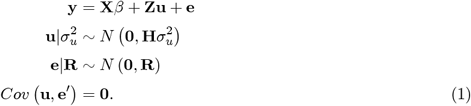

Above, *β* and **u** are unknown fixed effects and random breeding values, respectively; **X** and **Z** are known fixed incidence matrices and **e** is a vector of residuals with covariance matrix **R**, left unspecified at this point. **H** is a similarity or kernel matrix derived from both genomic and pedigree information (Misztal et al. 2009; Aguilar et al. 2010; Christensen and Lund 2010) and 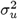 is the variance captured by **H**, and interpretable parametrically as the variance of breeding values. The ssGBLUP mixed model equations are

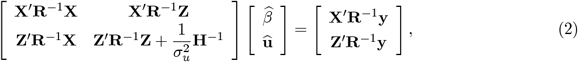

In (1) and (2), **H** stems from 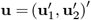, where **u**_1_ pertains to individuals in a pedigree without marker genotypes, and **u**_2_ to those in the pedigree but genotyped. Note that in SS all individuals are assumed to be pedigreed and the genealogical relatedness is conveyed by a known numerator relationship matrix (Henderson 1976) with form

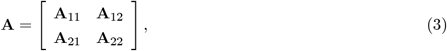

where **A**_11_ (**A**_22_) contains relationships among ungenotyped (genotyped) individuals and 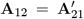 contains relationships between individuals in sets 1 and 2. The matrix **H** used in ssGBLUP has the (relatively) simple inverse

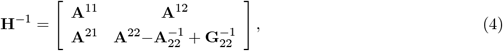

where **A**^*ij*^ is an appropriate partition of **A**^−1^, an inverse matrix derived from the pedigree using simple rules (Henderson 1976), and **G**_22_ is a full-rank genomic relationship matrix, e.g., Van Raden (2008). In animal breeding the order of **G**_22_ is typically much smaller than the order of **A**. Since **H**^−1^ is easy to obtain (assuming **G**_22_ and can **A**_22_ can be inverted), given **R** and 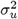, then 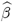 and 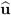 in (2) yield the (SS) best linear unbiased estimator of *β* and the (SS) best linear unbiased predictor (BLUP) of **u**, respectively. All individuals, genotyped or not, end up possessing an estimate of breeding value relative to the similarities conveyed by matrix **H**; if genotyping is abundant 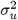 tends to 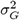 and in the absence of markers tends to 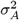. These two parameters give bounds for 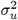. A stumbling block in solving (2) may be the calculation of 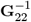 and 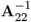, but see Misztal (2016), Masuda et al. (2016) and Strandén et al. (2017) for tricks. Individuals with pedigree but without phenotypes can be included by an appropriate expansion of **A**.

However, individuals that have markers but lack genealogies (set 3, say) do not have “an **A** to match”; **G** can be expanded, but the partitions **G**_33_, **G**_32_(**G**_23_) and **G**_31_(**G**_13_) lack counterparts in **A**. It is not obvious how the mixed model equations should be formed in such a case but techniques such as “unknown parent groups” can be used for such purpose. Technically, SS provides for imputations of genomic values from pedigree, but not the opposite.

Consider a Bayesian view of ssGBLUP. From theory, if a data generating process is **y** ~ *N* (**T***θ*, **D**) and the unknown location vector *θ* has prior distribution *θ* ~ *N* (*α*, **P**_*prior*_), the posterior is 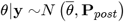, where

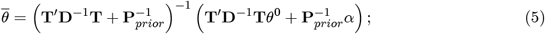

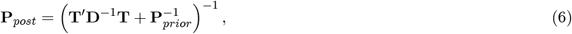

(e.g., Sorensen and Gianola, 2002). Above, *θ*^0^ is any solution to the possibly indeterminate “least-squares” system **T**^*^**D**^−1^**T***θ*^0^ = **T**′**D**^−1^**y** (Dempfle 1977). By definition, 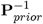 and 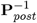 are the prior and posterior precision matrices, i.e., the inverses of the covariance matrices **P**_*prior*_ and **P**_*post*_, respectively. Note that the posterior expectation 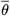 is a matrix-weighted average of *θ*^0^ and of the prior mean vector *α*, where the matrix weights are the “data precision” **T**′**D**^−1^**T** (coefficient or information matrix in least-squares) and the prior precision 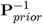. The posterior precision matrix is 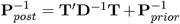, i.e., the sum of the prior precision plus the data precision. The Bayesian interpretation of ssGBLUP is implicit in derivations in Aguilar et al. (2010). In SS methodology, *θ* = (*β*′, **u**′)′; *β* is assigned a flat prior; 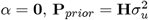 and 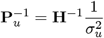 is the precision matrix of the prior distribution of **u**. Further, 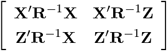 is the information from the observations (data precision) or from the Bayesian likelihood, equivalently.

## 3 Hy-BLUP

### 3.1 Setting

Let there be three disjoint classes of individuals: A) with phenotypes (**y**_1_) and pedigree but without marker genotypes (breeding values in vector **u**_1_); B) with phenotypes (**y**_2+_), pedigree and genotypes (breeding values in **u**_2+_), and C) with genotypes, with or without phenotypes but lacking pedigree (breeding values in **u**_2−_). Assuming **y**_2−_ has been observed in class (C), the linear model is

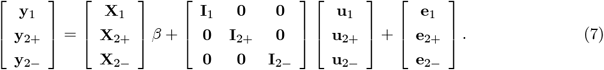

The identity matrices **I**. have orders equal to the number of individuals with phenotypes in each of the three classes. In a single-trait setting, assume

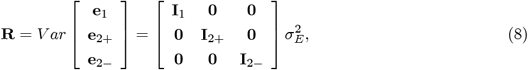

where 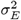 is a residual variance.

### 3.2 Precision matrices

As stated, in Bayesian analysis if an unknown vector has covariance matrix **K**, its precision matrix is **K**^−^, where **K**^−^ is a unique inverse provided the prior distribution is non-singular. If a vector of breeding values has full-rank covariance matrix 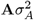 where 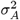 is a variance component, its precision matrix is therefore 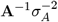. In the absence of genealogical information, individuals must be assumed to be genetically unrelated and independently distributed of other individuals in the population. All that is known is that breeding values have variance 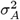, and distributed around a mean typically assumed null. For these individuals lacking genealogical information, one normatively takes 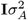 as prior covariance matrix and 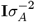 as joint precision. Likewise, if a vector of genomic (chip dependent) breeding values has covariance matrix 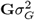, where 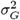 is a genomic variance (Lehermeier et al. 2017) and **G** is a genomic relationship matrix, the precision matrix is 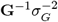; for individuals without genomic information, their joint precision is 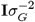. For location parameters *β* assigned improper priors (Bayesian counterparts of “fixed effects”, i.e., with infinite prior variance), the prior precision is a null matrix

For classes A, B, and C the precision matrix of the joint prior distribution of breeding values conveyed by pedigree information is

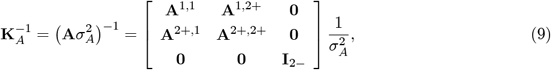

where **A**^*i,j*^ is an appropriate block of the inverse of the relationship matrix between animals with pedigree and **I**_2−_ has order equal to the number of individuals in class C. Similarly, the precision matrix of the prior distribution of genomic breeding values is

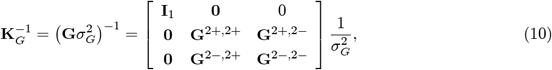

where

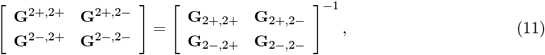

is the inverse of the genomic relationship matrix of all individuals genotyped.

Given 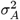 and 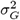, what is the prior precision of “true” breeding values conveyed by the two sources of information together, if assumed independent? The matter of independence is revisited in the discussion section of the paper. Reasoning as in Dempfle (1977), the joint prior precision matrix is the sum of the two precisions, that is

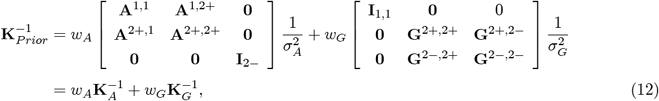

where *w*_*A*_ = *w*_*G*_ = 1 normatively. From a predictive perspective one may assigned distinct weights to pedigree and genome-based similarity matrices. For example, take *w*_*A*_ between 0 and 1 and *w*_*G*_ = 1 − *w*_*A*_, producing a weighted average of pedigree and genomic prior precisions. Here, when 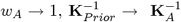. When 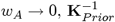 tends to 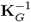.

If a flat prior (“infinite covariance matrix”) is assigned to *β*, the joint prior precision matrix of 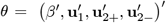 is

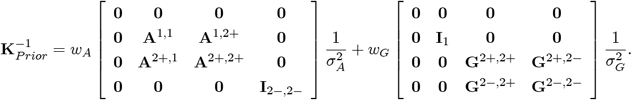

Now, the joint prior precision about *β* and **u** conveyed by the data 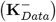 is given by the Fisher’s information matrix associated with the model, with all effects regarded as fixed. That is,

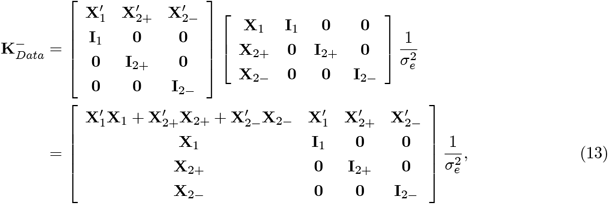

where 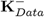 may be singular if at least one of the **X** matrices has incomplete column rank.

The residual variance in a model that uses pedigree information solely as in ABLUP 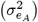, is expected to differ from that in GBLUP 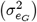. If estimates of residual variances and 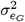, a residual variance suitable for Hy-BLUP could be

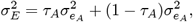

where

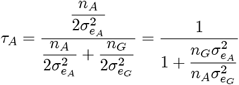

is a weight suggested by maximum likelihood asymptotic theory. The weight *τ*_*A*_ depends on the ratio between the number of animals with phenotypes and pedigree information (*n*_*A*_) and the number of animals with phenotypes and genomic information (*n*_*G*_) used to estimate the residual variances, and on the ratio 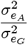.

### 3.3 Posterior expectations

In Bayesian Gaussian linear models with known variance components, the posterior expectation of *θ* satisfies a system of linear equations, as shown in (5). The coefficient matrix is the sum of the prior precision and of the precision conveyed by the data, and the right-hand sides are as in the mixed model equations of Henderson (1973) the if prior expectation of *θ* is null. For our model, the estimating equations can be written as

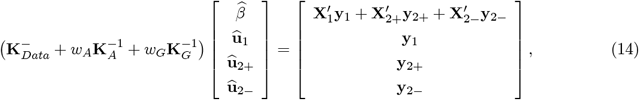

recalling that *w*_*A*_ = *w*_*G*_ = 1 are the “normative” (i.e., from Bayes theorem) weights. The coefficient matrix multiplied by 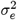 (recall that 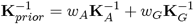) is

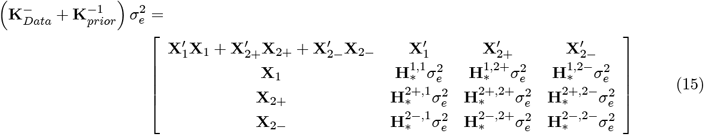

where

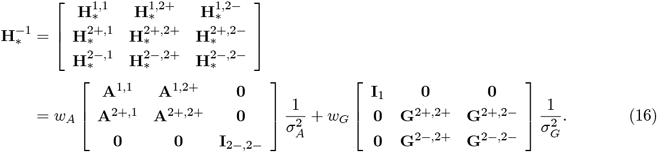

Forming a matrix such as 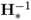 is standard in animal breeding and the computational complexities do not exceed those encountered in ssGBLUP. A special case is 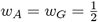 and 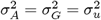. Here

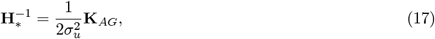

where

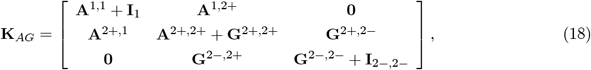

is a kernel associated with a random variable having variance 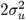. Clearly, the extent of shrinkage to 0 depends on the choice of weights.

Collecting terms, the mixed model equations (“hats” indicate posterior expectations) become

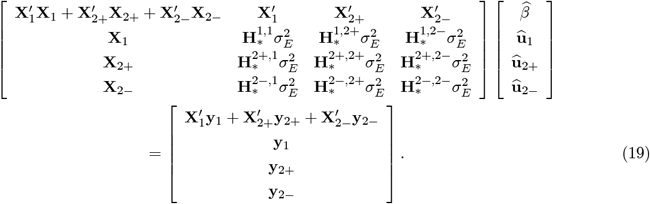

The posterior expectations have the usual frequentist interpretation: for the linear mixed model (7), 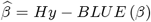, and 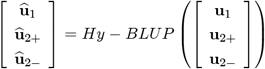, where *BLUE* and *BLUP* mean best linear unbiased estimator and best linear unbiased predictor, respectively, with respect to the covariance structure adopted. Calculation of *Hy* − *BLUE* and *Hy* − *BLUP* requires knowledge of 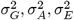 and an assignment of weights *w*_*A*_ and *w*_*G*_. With *w*_*A*_ = *w*_*G*_ = 1, the analysis yields the “normative” *Hy* − *BLUP* or *Hy* − *BLUP* (1, 1).

### 3.4 Special cases

Classes A and B comprise individuals recorded for some trait and fully pedigreed, but with only a fraction genotyped. Class C includes nominally unrelated individuals that have been genotyped but lack genealogy and records. Prediction of breeding values are sought for all members of classes A, B and C. Class C may comprise slaughtered meat animals that have been genotyped for some purpose but lack routine performance measurements, e.g., yearling weight in beef cattle. Because their genealogy is unknown, these animals cannot be traced to relatives in the population. However, there may be molecular similarity with animals in B, which possess records, thus enabling to obtain a prediction of class C breeding values for yearling weight.

A linear model for the observed yearling weight records is

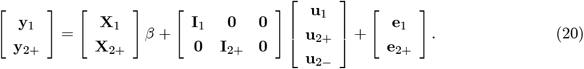

From (13) the precision contributed by the data is

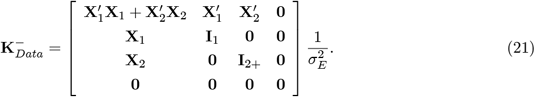

The data confer null precision to breeding values of individuals in class C because these do not have phenotypes. The mixed model equations for the example are

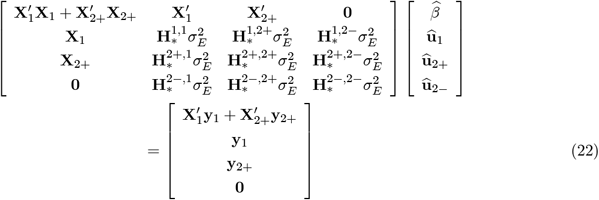

The 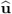 solutions of these equations give *Hy* − *BLUP* of breeding values for all animals, including those in class 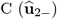.

### 3.5 Differences between ssGBLUP and Hy-BLUP

Toy examples in Supplementary File S1 illustrate how ABLUP, GBLUP, ssGBLUP and Hy-BLUP are calculated, giving different solutions. In ABLUP, GBLUP and ssGBLUP there is a single genetic variance (e.g., Aguilar et al. 2010; Christensen and Lund 2010; Fernando et al. 2014) whereas in Hy-BLUP there may be two: the infinitesimal and the genomic variances. Dispersion parameters are assumed known in all methods.

In ABLUP and GBLUP, the pedigree and genome-based similarity matrices are **A** and **G**, respectively. In ssGBLUP (Legarra et al. 2009; Fernando et al. 2014), the variance-covariance matrix of breeding values is

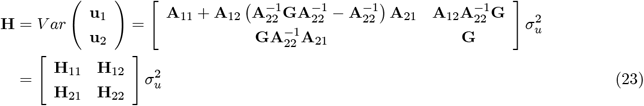

Here, the covariance between individuals possessing pedigree information only (i.e., 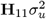) depends on similarities between individuals in classes A and B. The marginal distribution of **u**_2_, however, depends only on genomic similarities within class B.

BLUP is the multivariate regression of breeding values on phenotypes in a model without fixed effects. For ssGBLUP the regression of breeding values **u**_1_ on **y**_1_ is

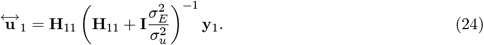

In ssGBLUP the regression function of **u**_1_ on **u**_2_ is 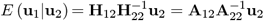, exactly as in a pedigree-based model. The regression function of **u**_2_ on **u**_1_ is

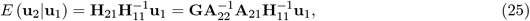

which it would be as in the infinitesimal model if **G** = **A**_22_, an event taking probability zero because **G** is a continuous-value random matrix with expectation **A** under idealized conditions (Van Raden 2008).

Since the variance-covariance matrix is the inverse of the precision matrix, it follows from (12) that for Hy-BLUP

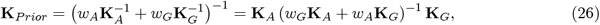

and

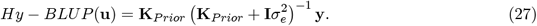

Insight about the form of **K**_*Prior*_ is obtained from consideration of a scalar situation. From (26) the variance of the prior distribution of an unknown breeding value is

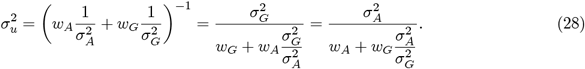

In *Hy* − *BLUP* (1, 1), *w*_*G*_ = *w*_*A*_ = 1 so 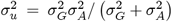. Expression (28), implies that the underlying breeding value modeled in Hy-BLUP has the same distribution as the random variable

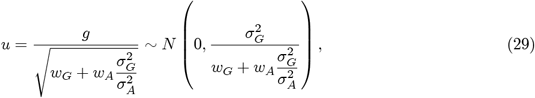

or as the random variable

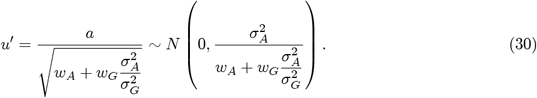

If 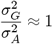 and *w*_*G*_ + *w*_*A*_ = 1, then *g, a*, and *u* (*u*′) have approximately the same distribution. From (29) and (30), if 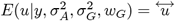, then

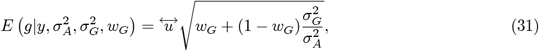

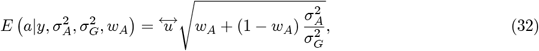

give the *Hy* − *BLUPs* of the “genomic and pedigree breeding value equivalents”, respectively.

More generally, the covariance matrix **K**_*Prior*_, implies that the vector of breeding values modeled in Hy-BLUP has the distribution

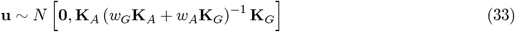

which would be **g** ~ *N* (0, **K**_*G*_) when *w*_*G*_ =1 and *w*_*A*_ = 0, or **a** ~ *N* (0, **K**_*A*_) when *w*_*A*_ =1 and *w*_*G*_ = 0.

## 4 Empirical evaluation of Hy-BLUP

### 4.1 Data

A wheat yield data set used in studies such as Gianola et al. (2016) and Gianola and Schön (2016) was downloaded from the BGLR package in `R` (Pérez and de los Campos 2014). The data comprises *n* = 599 inbred lines of wheat, each genotyped for *p* = 1279 binary markers that denote presence or absence of an allele at a locus. The phenotype was wheat grain yield per line planted in “environment 2”.

The lines are fully pedigreed and BGLR includes an additive relationship matrix among (**A**) of size 599 × 599 among them. A genomic relationship matrix (Van Raden 2008) among lines was built as **G** = **XX**′*/p* where matrix **X** had 599× 1279 (*p*) marker codes (0,1), with each column centered and standardized. Using all lines, we fitted the models **y** = **a**+**e** and **y** = **g**+**e**^*^ in pedigree-based and genome-enabled analyses, respectively, where **a** and **g** are pedigree and genomic breeding values, and **e** and **e**^*^ are model residuals. It was assumed that the vectors 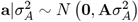 and 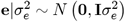 were mutually independent; the *σ*^2 ′^ *s* are variance components. For the genome-based analysis, 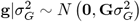 and 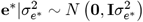 were taken as independent as well.

Maximum likelihood estimates of variances were obtained from the full data set and regarded as true values subsequently. BLUP of **a** and of **g** were calculated and denoted as 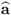 and 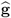, respectively. Under a Bayesian framework the posterior distributions of pedigree and genomic breeding values are 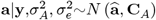 and 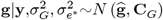, with variance components viewed as known hyper-parameters. The posterior expectations are

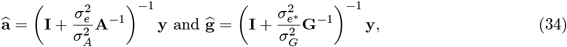

which are ABLUP and GBLUP, respectively, for the values of the variance parameters used. Further

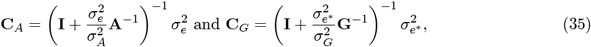

are posterior covariance matrices; in a frequentist setting, **C**_*a*_ and **C**_*g*_ are prediction error covariance matrices.

The maximum likelihood estimates of variance components were 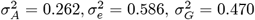 and 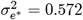; corresponding estimates of heritability were 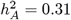 and 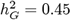. Using reciprocals of the asymptotic variances of the maximum likelihood estimates as weights, averages of genetic and residual variances were, 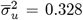 and 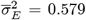, respectively. How variance component estimates were used for prediction is described in sections corresponding to each of four “Experiments” carried out.

### 4.2 Experiment 1: Hy-BLUP versus ABLUP and GBLUP

Experiment 1 had two objectives. First, contrasting Hy-BLUP (*w*_*A*_ = 0.25, 0.50, 0.75; *w*_*G*_ =1 − *w*_*A*_) and “normative” Hy-BLUP(*w*_*A*_ = *w*_*G*_ = 1) against ABLUP(*w*_*A*_ = 1, *w*_*G*_ = 0) and GBLUP (*w*_*A*_ = 0, *w*_*G*_ = 1). Second, evaluating the predictive ability of Hy-BLUP over a grid of weights. As mentioned, the target traits was wheat grain yield in environment 2.

All lines in the entire data set had phenotypes, pedigree and markers, and incompleteness of information was simulated by masking the appropriate type of data. Three designs were employed as described in Table 1. In all designs, each of the 599 lines was assigned at random to one of five groups of fixed sizes. Lines on groups 1, 3 and 4 were used for model training; groups 2 and 5 included testing set lines.

**Table 1.** Designs for evaluating Hy-BLUP (“Hybrid” BLUP). Groups 1, 3 and 4 used for training. Groups 2 and 5 (phenotypes masked) used as testing sets for evaluating predictive ability. Designs 1 and 2 employed constant variances. Design 3 assigned the same weights to residual variances as done to the corresponding similarity precision matrices, except for “normative” Hy-BLUP where 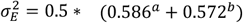. N_1_-N_5_ denote number of lines per group out of a total N=599.

| GROUP |  | Design 1 | Design 2 | Design 3 |
| --- | --- | --- | --- | --- |
| | **Weights placed on pedigree (w<sub>A</sub>) and genomic (w<sub>G</sub>) similarity** | $w_A=(0,0.25,0.50,0.75,1)$<br>$w_G=1-w_A \ (0 \leq w_G \leq 1)$<br>$w_G=w_A=1$ in<br>“normative” Hy-BLUP | $w_A=(0,0.25,0.50,0.75,1)$<br>$w_G=1-w_A \ (0 \leq w_G \leq 1)$<br>$w_G=w_A=1$ in<br>“normative” Hy-BLUP | $w_A=(0,0.25,0.50,0.75,1)$<br>$w_G=1-w_A \ (0 \leq w_G \leq 1)$<br>$w_G=w_A=1$ in<br>“normative” Hy-BLUP |
| | **Genetic and residual variances** | $\sigma_A^2 = 0.262$<br>$\sigma_G^2 = 0.470$<br>$\sigma_E^2 = \frac{1}{2} * 0.586^a$<br>$+ \frac{1}{2} * 0.572^b$ | $\sigma_A^2 = 0.262$<br>$\sigma_G^2 = 0.470$<br>$\sigma_E^2 = \frac{1}{2} * 0.586^a$<br>$+ \frac{1}{2} * 0.572^b$ | $\sigma_A^2 = 0.262$<br>$\sigma_G^2 = 0.470$<br>$\sigma_E^2 = w_A * 0.586^a$<br>$+ w_G * 0.572^b$ |
| 1.Pedigree and phenotypes | Training | N<sub>1</sub>=200 | N<sub>1</sub>=100 | N<sub>1</sub>=100 |
| 2.Pedigree | Testing | N<sub>2</sub>=25 | N<sub>2</sub>=150 | N<sub>2</sub>=150 |
| 3.Pedigree, markers and phenotypes | Training | N<sub>3</sub>=200 | N<sub>3</sub>=99 | N<sub>3</sub>=99 |
| 4. Markers and phenotypes | Training | N<sub>4</sub>=100 | N<sub>4</sub>=100 | N<sub>4</sub>=100 |
| 5.Markers | Testing | N<sub>5</sub>=74 | N<sub>5</sub>=150 | N<sub>5</sub>=150 |
|  |  | **N=599** | **N=599** | **N=599** |

Design 1 had 500 lines with phenotypes (groups 1, 3 and 4) and 99 without phenotypes (groups 2 and 5 with 25 and 75 lines, respectively). Designs 2 and 3 had 299 lines in groups 1, 3 and 4, and 300 lines in groups 2 and 5. Within lines having phenotypes, group 1 had pedigree only; group 3 had pedigree and markers, and group 4 had markers only. In lines without phenotypes, group 2 had pedigree information but no markers, and Group 5 had markers but lacked genealogy.

For the purpose of evaluating predictive ability in Experiment 1, 20000 bootstrap samples were taken in each of the designs. For example, in Design 1, 500 lines were used for model training and 99 lines without for testing predictions. The 99 pairs of prediction-masked phenotype values were re-sampled with replacement 20000 times for each of the five values of *w*_*A*_ in the grid and for “normative” Hy-BLUP (with *w*_*A*_ = *w*_*G*_ = 1). Likewise, in Designs 2 and 3, training and testing sets had 299 and 300 members, respectively; the 300 pairs of predictions-masked phenotype values were re-sampled 20000 times for each model fitted. Bootstrap samples were used to estimate the distributions of prediction metrics such as correlations and mean-squared errors.

As indicated in Table 1, genetic variances 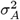 and 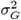 were kept constant across designs but there were differences in values of the residual variance components. In Designs 1 and 2, the residual variance was 0.579, the unweighted average of 0.586 and 0.572 from pedigree and genome-based analyses. In Design 3, the residual variance for “normative” Hy-BLUP was 0.579 while for other forms of Hy-BLUP the weights assigned to pedigree and genome-based precisions 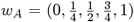, with *w*_*G*_ =1 − *w*_*A*_, were used for the residual variances. When *w*_*A*_ = 0, the analysis is driven entirely by genomic similarity, so *Hy* −*BLUP* yields GBLUP for the choice of variances indicated in Table 1; when *w*_*A*_ = 1, similarity derives entirely from pedigree and *Hy* − *BLUP* produces ABLUP.

By comparing estimated breeding values of lines in groups 2 and 5 with their realized phenotypes (masked in model training), estimates of realized prediction error and measures of agreement between predictions and predictands were obtained.

We illustrate how Hy-BLUP was implemented under Design 1. The training model for phenotypic values (subscripts denote groups) was

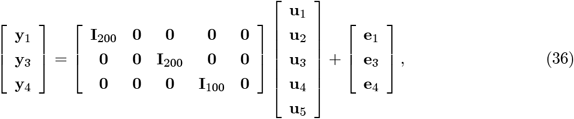

where a **0** denotes a null matrix of appropriate order. For *w*_*A*_ = 0.25, say, the precision of the prior distribution of breeding values is

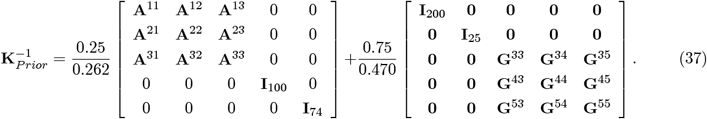

The precision matrix of the data, using (as an example) 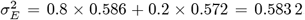, so 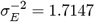, was

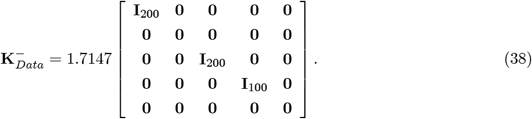

The estimated breeding values are obtained by solving equations (19) adapted to model (36). The same procedure was used for each value of *w*_*A*_ in the grid.

Figure 1 displays fitted values of linear regressions of phenotypes on estimated HY-BLUP breeding values of the *n* = 599 lines for some of the models fitted to the entire data set. Regressions for models with *w*_*A*_ = 0.25 (*w*_*G*_ = 0.75) and *w*_*A*_ = 0.75 (*w*_*G*_ = 0.25) are not shown to reduce clutter. For ABLUP (*w*_*A*_ = 1, *w*_*G*_ = 0) and GBLUP (*w*_*A*_ = 0, *w*_*G*_ = 1), variance components were from separate pedigree and genome based analyses, respectively. In other HY-BLUP fits, the residual variance was the average of the two estimates of residual variance; “normative” Hy-BLUP had *w*_*A*_ = *w*_*G*_ = 1. The largest *R*^2^ (0.72) was for ABLUP, followed by Hy-BLUP 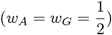 with *R*^2^= 0.64, GBLUP (0.62) and “normative” Hy-BLUP (0.58); for *w*_*A*_ = 0.25, *w*_*G*_ = 0.75 and *w*_*A*_ = 0.75, *w*_*G*_ = 0.25, the *R*^2^ values were 0.63 and 0.67, respectively. The steepest line was for “normative” HY-BLUP. This specification corresponds to the sharpest state of prior information considered, i.e., with the lowest prior variance and less dispersion among estimated breeding values (x-axis in Figure 1), thus yielding a larger slope.

**FIGURE 1.**
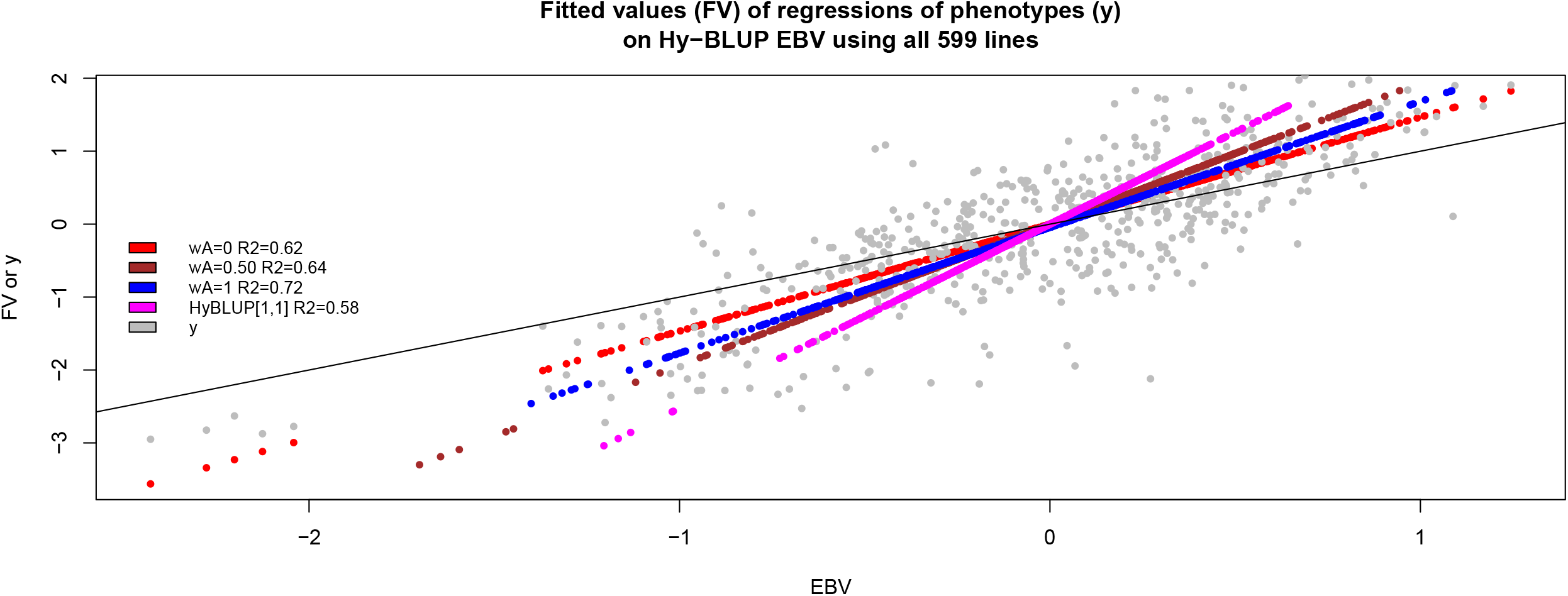
Fitted values of linear regressions of phenotypes (y, wheat grain yield) on Hy-BLUP estimated breeding values (EBV). *w*_*A*_ *= weight on pedigree. R*^2^ *= coefficient of determination.* Hy-BLUP [1,1]: normative weights.

As an example, take 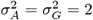 and 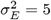. The prior precision for both ABLUP or GBLUP is 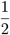 and the variance ratio is 5*/*2= 2.5. For Hy-BLUP 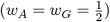, the prior precision is 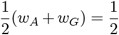 and the variance ratio is also 2.5. When *w*_*A*_ = *w*_*G*_ =1 (“normative” Hy-BLUP), the prior precision is 1 and the variance ratio is equal to 5, producing more shrinkage. In another example, take 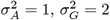 and 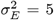. The prior precisions for ABLUP and GBLUP are 1 and 1*/*2, and the respective variance ratios are 5 and 2.5. For weights between 0 and 1, the largest possible precision is with *w*_*A*_ =1 (ABLUP). However, for Hy-BLUP(1,1), the prior precision is 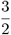 and the variance ratio takes the value 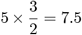. These two examples illustrate why Hy-BLUP yields point estimates with smaller absolute values than any Hy-BLUP with positive weights *w*_*A*_ =1 − *w*_*G*_ and *w*_*A*_ + *w*_*G*_ = 1.

Figures 2 and 3 depict pair-wise plots of Hy-BLUP estimates of breeding values obtained with *w*_*A*_ =1 (ABLUP), 0.75, 0.50 and 0.25 and *w*_*A*_ = 0 (GBLUP) for a random realization of Design 1 in Table 1; EBVs from Hy-BLUP(1,1) are shown also. The left panel illustrates that, with GBLUP (*w*_*A*_ = 0), the 74 null estimates of breeding value obtained with ABLUP (*wA* = 1) were disambiguated; Hy-BLUP(*w*_*A*_ =0.75,0.50, 0.25, 0; *w*_*G*_ = 1−*w*_*A*_) and Hy-BLUP(1,1) produced a single EBV=0, corresponding to a line whose centered phenotypic value was 0. The scatterplots in the four panels in Figures 2 and 3 show that the absolute values of Hy-BLUP(1,1) estimates were smaller than those from other specifications of weights. Correlations between EBVs were such that the lowest one (0.79) was between ABLUP and GBLUP. Hy-BLUP(1,1) had a stronger correlation with EBVs from *w*_*A*_ = 0.25, 0.50 and 0.75 (0.98-0.99) than with those found with *w*_*A*_ =0 (GBLUP) or *w*_*A*_ =1 (0.92).

**FIGURE 2.**
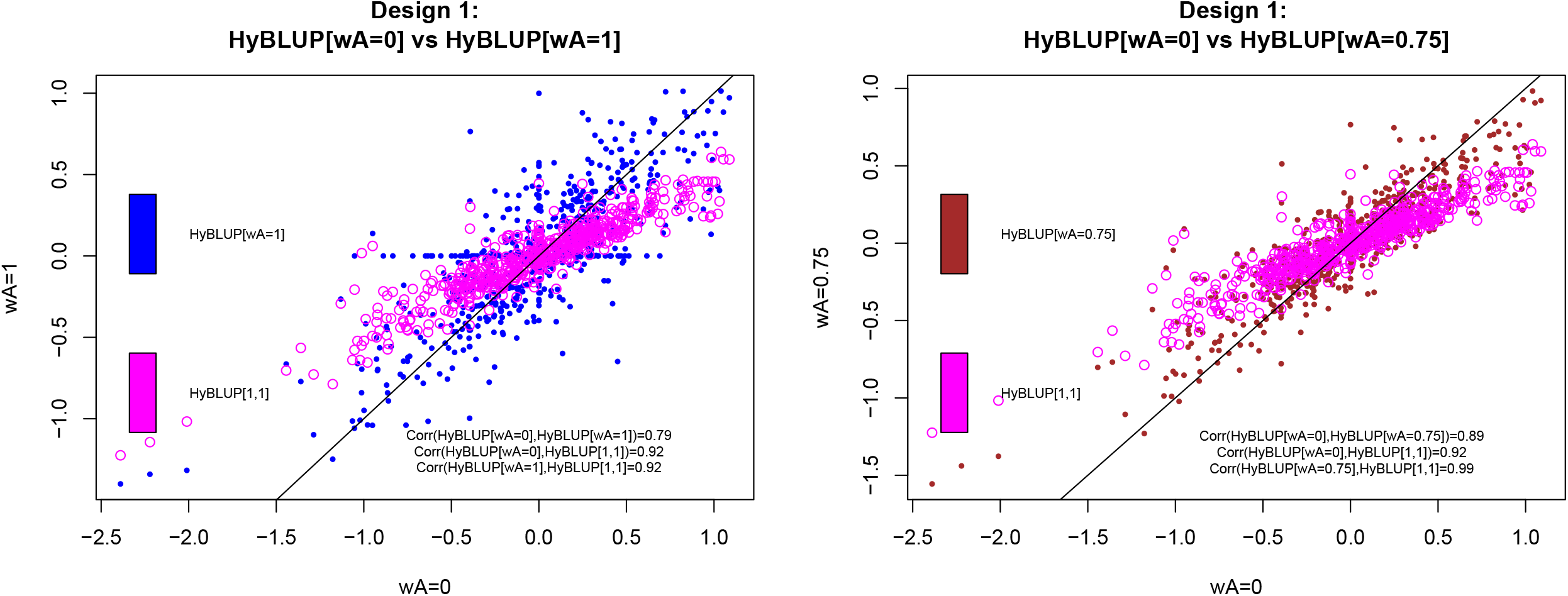
Experiment 1-Design 1. Scatter plots of estimated breeding values obtained with Hy-BLUP. *w*_*A*_ *= weight on pedigree:* 0, 0.75 and 1. Hy-BLUP [1,1]: normative weights.

**FIGURE 3.**
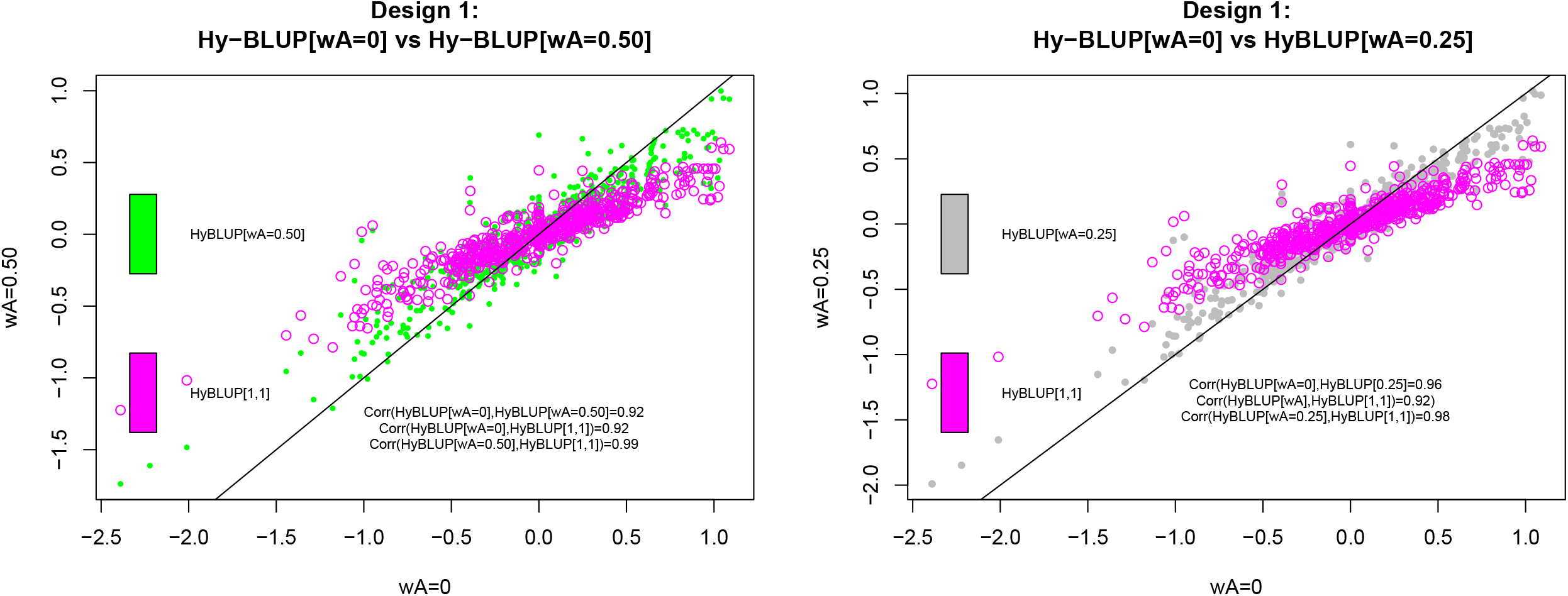
Experiment 1-Design 1. Scatter plots of estimated breeding values obtained with Hy-BLUP. *w*_*A*_ *= weight on pedigree*: 0, 0.25 and 0.50. Hy-BLUP [1,1]: normative weights.

Densities of predictive correlations and mean squared errors estimated from 20000 bootstrap samples obtained from a realization of Design 1 are in Figure 4. Results for testing sets represented by groups 2 (pedigree only) and 5 (markers only) are shown in the left and right-side panels, respectively. As indicated by the relatively flat densities, uncertainty was large. When test set individuals had pedigree information only, the median predictive correlation ranged from 0.22 for Hy-BLUP(1,1) to 0.28 for Hy-BLUP(*w*_*A*_ = 0); in group 5 (panel at right) predictive ability was nil. Densities overlapped, without clear differences between the various levels of *w*_*A*_ as indicated by the medians of the distributions. Densities for MSE were also flat and differences among methods were not marked. When individuals in the testing set had pedigree information only (bottom left panel), median MSE differed by little across the various specifications of weights on pedigree, with a slight advantage for *w*_*A*_ = 1 (ABLUP). When the test set individuals had only markers (bottom right panel), MSE was lowest for Hy-BLUP(1,1) and highest for *w*_*A*_ = 1. Mean squared error was smaller when test set individuals had marker information only (bottom right panel, Figure 4) than when they had pedigree information only (bottom left panel).

**FIGURE 4.**
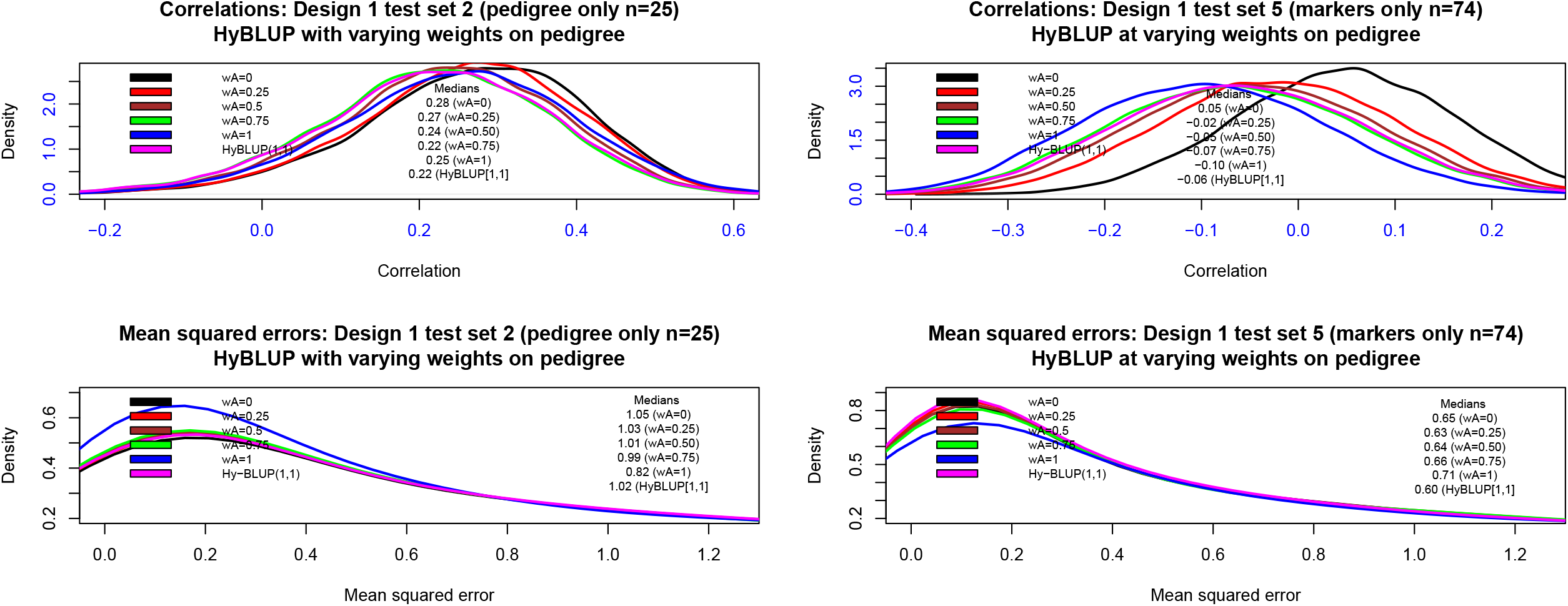
Experiment 1-Design 1. Predictive correlations and mean-squared errors for various implementations of Hy-BLUP using 20,000 bootstrap samples. *w*_*A*_ *= weight on pedigree*: 0, 0.25, 0.50, 0.75, 0.75,1. Hy-BLUP [1,1]: normative weights.

Predictive densities for Design 2 are shown in Figure 5. The densities for the correlations were not sharp (large uncertainty) although symmetric; the value 0 appeared at high density so differences among methods were not detected. Predictive mean squared error was smaller for *w*_*A*_ = 1 than for other specifications of weights and the bootstrap-based densities suggested smaller prediction errors when test-set individuals had markers only (bottom right panel).

**FIGURE 5.**
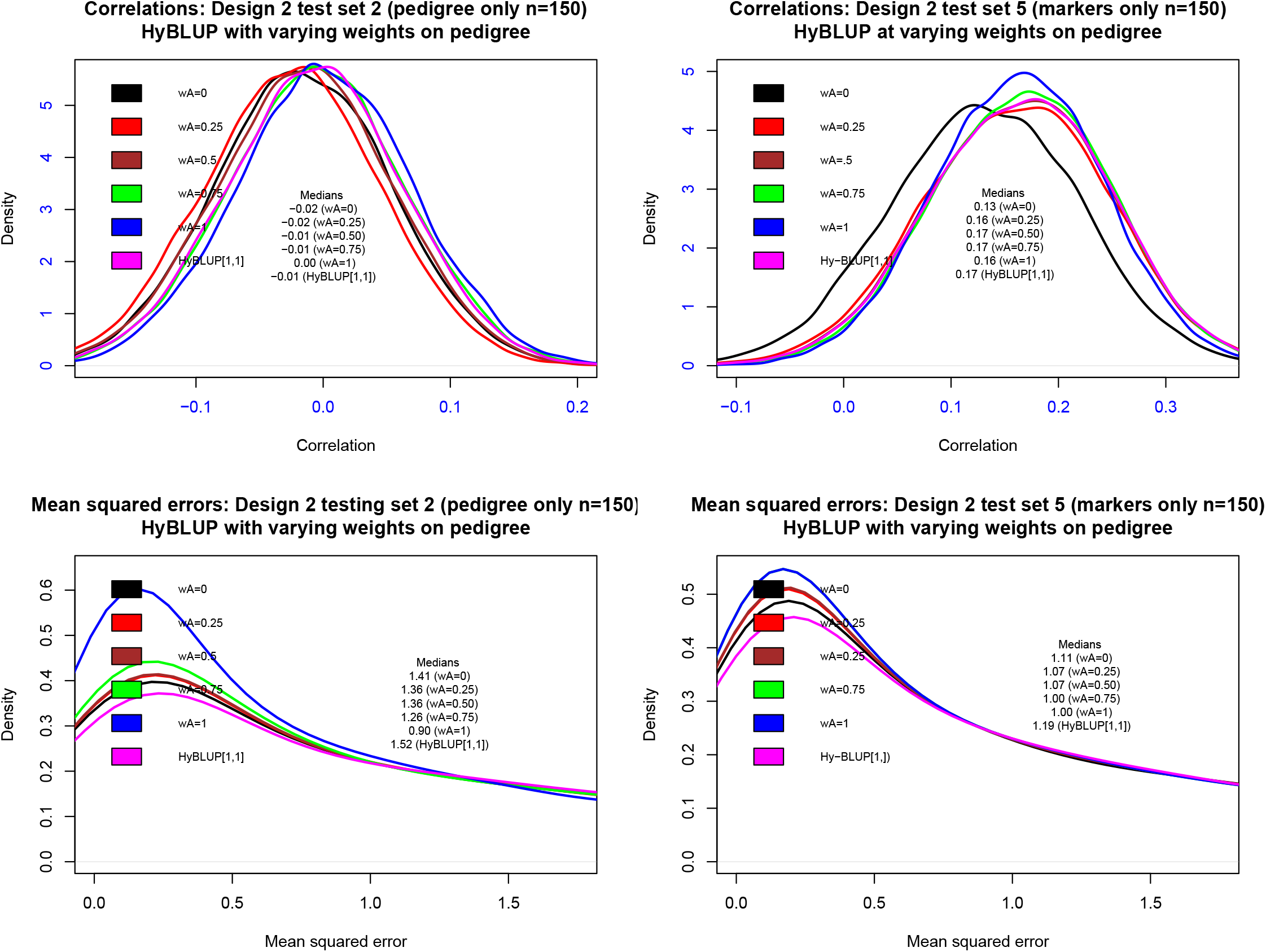
Experiment 1-Design 2. Predictive correlations and mean-squared errors for various implementations of Hy-BLUP using 1500 random replications. *w*_*A*_ *= weight on pedigree*: 0, 0.25, 0.50, 0.75, 0.75,1. Hy-BLUP [1,1]: normative weights.

Recall (Table 1) that in Design 3, Hy-BLUP implementations with *w*_*A*_ = (0, 0.25, 0.50, 0.77, 1) and *w*_*G*_ = 1 − *w*_*A*_ also assigned the same weights when averaging residual variances from separate genome and pedigree-based analysis, while the normative Hy-BLUP used the simple average. As shown in Figure 6, there was overlap between the six different densities, the same holding true when individuals in test sets had only genealogical (left panels) or genomic data. Predictive correlations were somewhat larger when test set individuals had markers only than when they had pedigree information. In group 2 (left panel) median correlations ranged between 0.00 to 0.07, and in group 5 (right panel) they ranged between 0.08 and 0.11. There was a slight tendency for predictive mean-squared errors to be smaller when genomic and pedigree precision matrices were blended (i.e., when placing non-null weights on precision matrices) than for the “pure” situations *w*_*A*_ = 0 and *w*_*G*_ = 1. Among the various specifications of Hy-BLUP the “normative” model had the smallest average squared prediction error.

**FIGURE 6.**
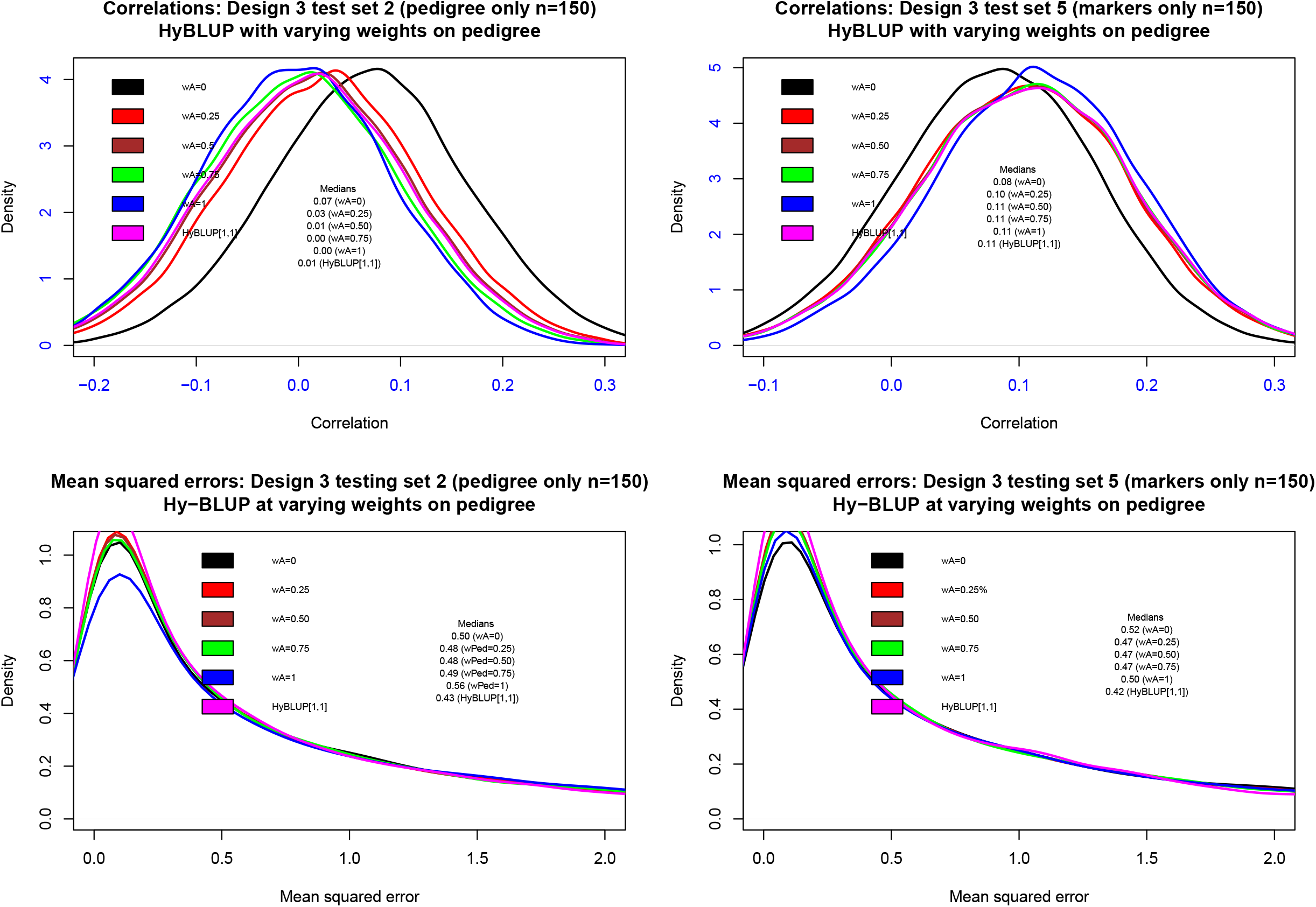
Experiment 1-Design 3. Predictive correlations and mean-squared errors for various implementations of Hy-BLUP using 1500 random replications. *w*_*A*_ *= weight on pedigree*: 0, 0.25, 0.50, 0.75, 0.75,1. Hy-BLUP [1,1]: normative weights.

Overall, Experiment 1 provided a proof of concept that Hy-BLUP can accommodate complex data patterns induced by variation in type and amount of input information in training and testing set individuals, without hampering and perhaps improving the ability of predicting yet-to be-observed phenotypes.

### 4.3 Experiment 2: Hy-BLUP versus ssGBLUP

In Experiment 2, Hy-BLUP was compared with ssGBLUP. A training-testing design was adopted. All individuals in training sets had phenotypes and pedigrees but some lacked marker information, which is the situation addressed by ssGBLUP. Individuals in testing sets had both markers and pedigrees. Table 2 describes the compositions of training-testing sets in six layouts, each of which was randomly replicated 1500 times. Training sets had from 200 to 400 lines with phenotypes, and testing sets had from 199 to 399 lines. Empirical studies with a fixed total sample size (e.g., Erbe et al. 2010; Gianola and Schön 2016) have indicated that when testing sets are made larger at the expense of training set sizes, predictive correlations tend to be weaker but less variable. Conversely, when training sets are made larger at the expense of testing sets, predictions are better but less precise. On this basis, one could expect layouts 4, 5 and 6 to deliver larger predictive correlations and larger mean-squared errors than layouts 1, 2 and 3.

Two versions of Hy-BLUP were run: 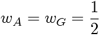 and *w* _*A*_ = *w*_*ZG*_ = 1. The “implicit genetic variance” was 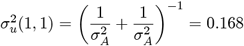 and 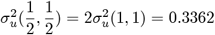. as suggested by formula (26). ssGBLUP was computed as described in the Excursus section of the paper; the genetic variance here was 0.328 (average of pedigree and genome-based maximum likelihood estimates using reciprocals of asymptotic variances as weights) and 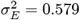, same as for Hy-BLUP.

**Table 2.** Layouts for comparing Hy-BLUP (“Hybrid” BLUP) versus SS-BLUP (single-step BLUP). Within a layout, 599 inbred wheat lines were assigned at random to training (Groups 1 and 2) and testing (Group 3) sets. Individuals in the training set had phenotypes and pedigree, but some lacked marker information. Members of testing sets had pedigree and marker data and phenotypes were masked. The structure of the six layouts was fixed and there were 1500 random replications of the allocation.

|  | **Layout** | **1** | **2** | **3** | **4** | **5** | **6** |
| --- | --- | --- | --- | --- | --- | --- | --- |
| **Group** | **Type of information** | No. of lines | No. of lines | No. of lines | No. of lines | No. of lines | No. of lines |
| 1. Training | Pedigree and phenotypes | 100 | 100 | 200 | 200 | 300 | 100 |
| 2. Training | Pedigree, markers and phenotypes | 100 | 200 | 100 | 200 | 100 | 300 |
| 3. Testing | Pedigree and markers | 399 | 299 | 299 | 199 | 199 | 199 |
|  | **Total No. of lines** | **599** | **599** | **599** | **599** | **599** | **599** |

Shrinkage is controlled mainly by the ratios of residual to genetic variances. These were 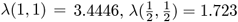 and *λ*_*SS*_ = 1.767 for the two sets of weights (Hy-BLUP) and SS, respectively. The variance ratios give an approximate idea of the extent of shrinkage, because the expressions are based on a scalar version of (26) while the effective shrinkage also involves the off-diagonals of the similarity matrices. For each layout and random repetition, Hy-BLUP and ssGBLUP were computed and predictions of phenotypes in the testing set were compared against realized values (masked during training). Quality of prediction was gauged using correlation and mean absolute and squared errors.

Hy-BLUP and ssGBLUP predictions were closely correlated. Based on 9000 runs (1500 replications for each of the six layouts), correlations between ssGBLUP and either 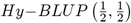 or *Hy*−*BLUP* (1, 1) ranged between 0.84 and 0.98. Correlations between the two versions of Hy-BLUP ranged between 0.993 and 0.999, so the two methods were almost perfectly associated. Correlations between ssGBLUP and *Hy* − *BLUP* (1, 1) were 0.90, 0.93, 0.94, 0.95, 0.95, and 0.95 in layouts 1-6, respectively.

Table 3 presents median, minimum and maximum values of predictive correlations (PCOR) and mean-squared error (MSE) over the 1500 random reconstructions of each of the layouts in Table 2. Predictive correlations were low and variable, and 0 was assigned a high density (curves not presented). The largest median PCOR was observed for ssGBLUP in 2 out of the six layouts and for Hy-BLUP(1,1) in 4 layouts; however, differences were not meaningful. On the other hand, MSE was lowest for *Hy* − *BLUP* (1, 1) in all replicates, at about 88-91% of that observed for ssGBLUP. Differences in PCOR or MSE between 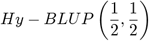 and *Hy* − *BLUP* (1, 1) were negligible.

**Table 3.** Predictive correlations (PCOR) and mean-squared errors (MSE) for two implementations of Hy-BLUP (“Hybrid” BLUP), Hy-BLUP(0.5,0.5) and HY-BLUP(1,1), and SS-BLUP (“single-step BLUP). The structure of the six layouts is described in Table 2. The largest median PCOR and smallest MSE for a given layout are boldfaced. Statistics are based on 1500 random replications of each layout.

|  |  | PCOR |  |  | MSE |  |  |
| --- | --- | --- | --- | --- | --- | --- | --- |
| Layout |  | SS-BLUP | Hy-BLUP<br>(0.5,0.5) | Hy-BLUP<br>(1,1) | SS-BLUP | HyBLUP<br>(0.5,0.5) | Hy-BLUP<br>(1,1) |
| **1** | Median | **0.040** | 0.038 | 0.038 | 1.110 | 1.043 | **1.013** |
|  | Minimum | -0.116 | -0.112 | -0.114 | 0.941 | 0.897 | 0.866 |
|  | Maximum | 0.209 | 0.222 | 0.222 | 1.301 | 1.205 | 1.171 |
| **2** | Median | **0.030** | 0.027 | 0.027 | 1.148 | 1.074 | **1.027** |
|  | Minimum | -0.177 | -0.181 | -0.181 | 0.885 | 0.844 | 0.799 |
|  | Maximum | 0.212 | 0.207 | 0.202 | 1.366 | 1.290 | 1.234 |
| **3** | Median | 0.027 | 0.027 | **0.028** | 1.148 | 1.073 | **1.024** |
|  | Minimum | -0.173 | -0.166 | -0.166 | 0.929 | 0.866 | 0.819 |
|  | Maximum | 0.216 | 0.218 | 0.222 | 1.433 | 1.335 | 1.260 |
| **4** | Median | 0.000 | 0.005 | **0.006** | 1.194 | 1.110 | **1.046** |
|  | Minimum | -0.228 | -0.236 | -0.232 | 0.837 | 0.803 | 0.769 |
|  | Maximum | 0.259 | 0.289 | 0.289 | 1.574 | 1.456 | 1.353 |
| **5** | Median | 0.004 | 0.009 | **0.011** | 1.191 | 1.108 | **1.044** |
|  | Minimum | -0.221 | -0.211 | -0.205 | 0.866 | 0.806 | 0.773 |
|  | Maximum | 0.216 | 0.210 | 0.222 | 1.575 | 1.457 | 1.359 |
| **6** | Median | 0.004 | 0.011 | **0.012** | 1.199 | 1.108 | **1.053** |
|  | Minimum | -0.206 | -0.228 | -0.218 | 0.883 | 0.806 | 0.778 |
|  | Maximum | 0.214 | 0.234 | 0.225 | 1.597 | 1.457 | 1.379 |

Figure 7 displays pairwise plots of realized mean absolute prediction errors (1500 values per layout) between *ssGBLUP* and *Hy* −*BLUP* (1, 1). Absolute prediction errors were highly correlated as indicated by the scatterplots. However, *Hy* − *BLUP* (1, 1) exhibited smaller errors systematically; the median absolute prediction error was about 4-7% smaller than that of *ssGBLUP* over the six layouts.

**FIGURE 7.**
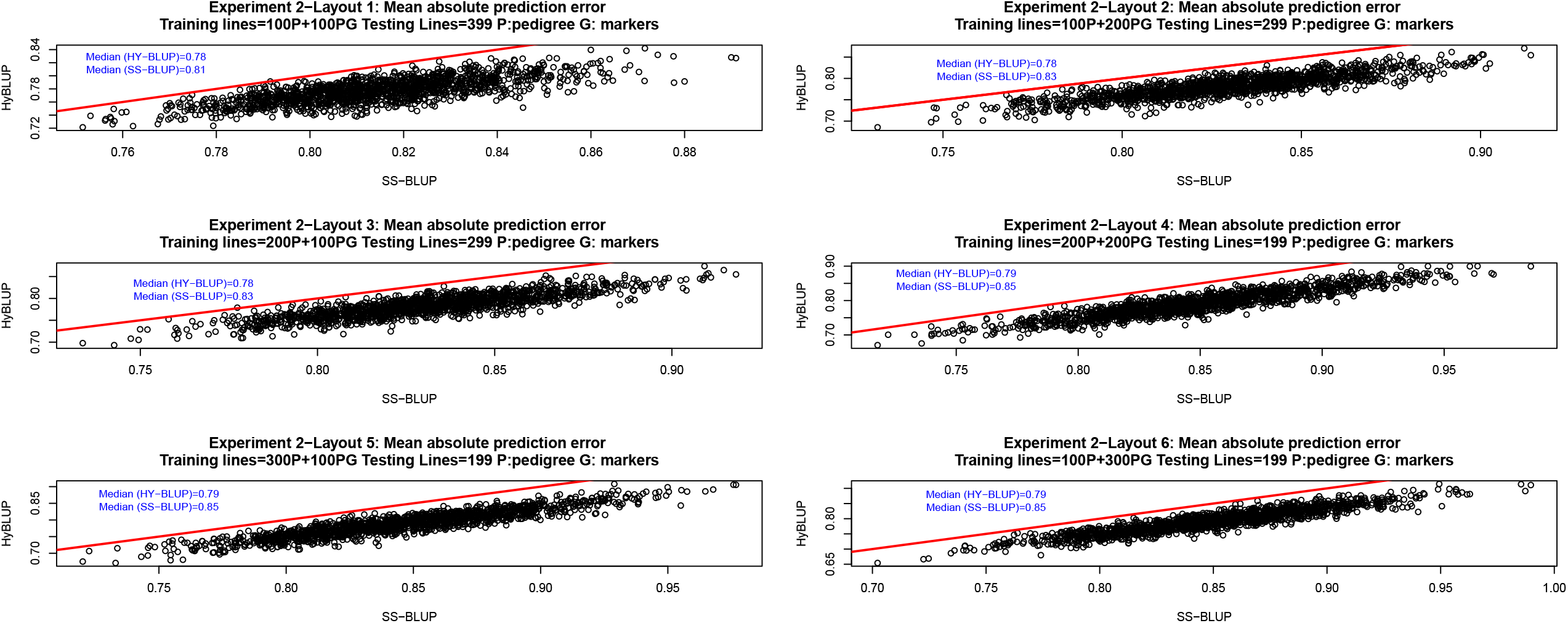
Experiment 1-Layouts 1-6. Mean absolute prediction error from 1500 replications: Hy-BLUP [1,1] versus single-step genomic BLUP (SS-BLUP).

Overall, Experiment 2 indicated that the predictive ability of ssGBLUP and *Hy* − *BLUP* was weak (Table 3). Although predictions were strongly aligned in the correlation sense, *Hy* − *BLUP* was more accurate, i.e., prediction errors were smaller. A plausible theoretical explanation is that *Hy* − *BLUP* is a method in which precision matrices are blended or “averaged” in some sense while *ssGBLUP* uses a form of imputation (e.g., Aguilar et al. 2010). *Hy* − *BLUP* shares conceptual commonality with the notion of model averaging or “kernel averaging (Hoeting et al. 1997; Sorensen and Gianola 2002; de los Campos et al. 2010; Tusell et al. 2014; Gianola and Schön 2016)

### 4.4 Experiments 3 and 4: Hy-BLUP versus ssGBLUP with estimated variances and unequal weights

Using fixed values of the variance components, as in experiment 2, may have unintentionally favored one of the two methods compared. The design shown in Table 2 was kept in Experiment 3 and genetic and residual variance components were estimated, by method and replication, taking 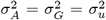 in Hy-BLUP. Each of the six layouts was randomly replicated 2000 times so there were 12000 sets of estimates of variances of breeding values and of residuals by method. The **H** (ssGBLUP) and **K**_*Prior*_ (Hy-BLUP) similarity matrices given in (4) and (12), respectively, were reconstructed in each realization. For building **K**_*Prior*_, the weights were 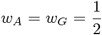. An eigen-decomposition of **H** and of **K**_*Prior*_, together with the *R* function *optim* were used for computing maximum likelihood estimates. Predictive correlations, mean absolute errors and mean-squared errors were calculated in the testing sets with sizes as indicated in Table 2.

Overall the 12000 runs, median residual variances were 0.51 (0.43-0.59) for 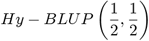 and 0.58 (0.48-0.68) for ssGBLUP; median estimates by layout were 0.51-0.52 and 0.58 for the two methods, respectively. Heritability was defined as the fraction of the total variance captured by the similarity (kernel) matrices **H** and **K**_*Prior*_. Heritability has a clear parametric meaning when the kernels are **A** and **G**, yielding the “infinitesimal” and “genomic” heritabilities, respectively (de los Campos et al. 2015). The estimate in ssGBLUP is a function of the fraction of individuals that have markers; as the fraction tends to 1, ssGBLUP heritability tends to genomic heritability. In *Hy* − *BLUP*, which is a blending or averaging procedure, the heritability parameter plays a tuning role, controlling the extent of shrinkage exerted by the prediction machine. Figure 8 depicts the 12000 estimates of heritability for Hy-BLUP (left panel) and ssGBLUP (right panel), colored by layout. The white line represents fitted values from a loess regression of heritability estimates on sample number (1:12000) ordered sequentially from layouts 1 through 6. The local polynomial fitted was of second order and the span was 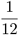, representing half the fraction of the number of samples in a given replicate relative to the total. The local regression flags “regions” sharing common values. For example, estimates in layouts 1 and 2 were higher than in layouts 3 and 4 but similar to those in layout 6. However, differences were minor: in layouts 1 and 2 median heritability was 0.58 for 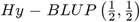 and 0.42 in ssGBLUP; in layouts 3 and 4 the respective values were 0.56 and 0.40. Regularization (shrinkage to 0) was stronger in ssGBLUP than in Hy-BLUP because of larger ratios of residual to kernel variances. Median values of 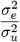 were 1.44 (0.87-2.62) and 0.76 (0.54-1.14) in ssGBLUP and 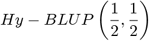, respectively.

**FIGURE 8.**
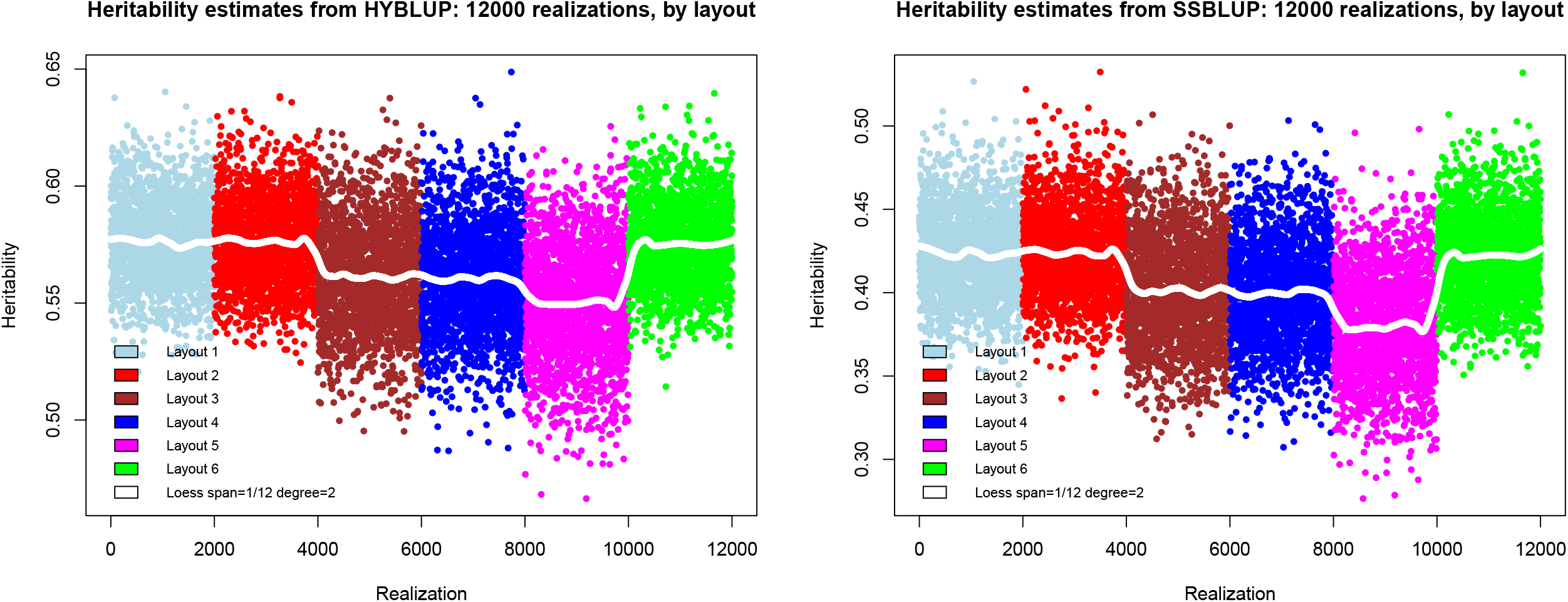
Heritability estimates from Hy-BLUP and single-step genomic BLUP(SS-BLUP). There are six layouts with 2000 random replications per layout. The white line represents fitted values from a local regression of estimate on sample sequence value (see text).

Figure 9 displays the 12000 estimates of differences (HyBLUP-ssGBLUP) in predictive correlations (left panel) and mean absolute prediction error (right panel). Differences in predictive correlations between procedures were not detected in any of the layouts and the curve of loess fitted valued was essentially flat. 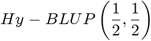 had slightly smaller mean absolute and squared prediction errors than ssGBLUP in layouts 1, 2 and 3, i.e., in those with larger (smaller) testing (training) sets. Overall, experiment 3 suggested that Hy-BLUP delivered predictions of similar or slightly better quality than ssGBLUP when regularization parameters were estimated from training samples. In Design 2, with fixed values of variance components, ssGBLUP exhibited systematically larger absolute prediction errors.

**FIGURE 9.**
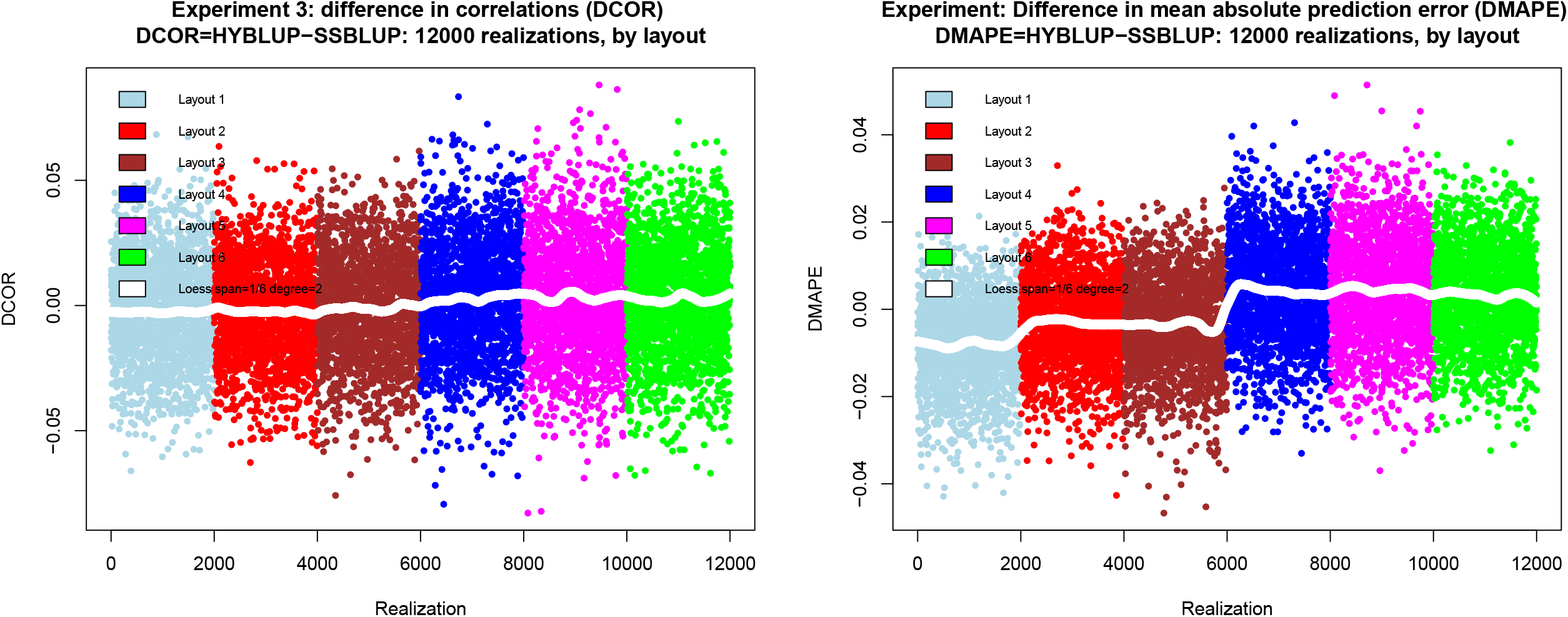
Difference in predictive correlation and mean absolute prediction error between Hy-BLUP and single-step genomic BLUP(SS-BLUP). There are six layouts with 2000 random replications per layout. The white line represents fitted values from a local regression of estimate on sample sequence value (see text).

In Experiment 4, we examined the impact of varying weights, with re-estimation of variance components as in Experiment 3, on out-of-sample correlation and mean absolute error of prediction. The six layouts in Table 2 were maintained and the weights assigned to precision from pedigree were *w*_*A*_ = 0.25, *w*_*A*_ = 0.50 and *w*_*A*_ = 0.75, *w*_*G*_ = 1 − *w*_*A*_. Each layout-weight combination was replicated 2000 times, completely at random. In short (results not shown), larger prediction errors were associated with a lower predictive correlation, as expected. Absolute values of differences between Hy-BLUP and ssGBLUP did not exceed 0.004 and 0.008 for predictive correlations and mean absolute error of prediction, respectively. Patterns were similar across the three values of *w*_*A*_.

Experiments 3 and 4, employing variances estimated from training samples indicated that the differences between the two prediction methods were negligible from the predictive correlation sense but Hy-BLUP was somewhat more accurate in terms of smaller prediction errors.

## 5 Discussion

### Focus

We presented a method for prediction of complex traits (Hy-BLUP) that combines genealogical and molecular marker information. The procedure is suited to situations in which some individuals have both types of information while others have pedigree or genomic data only. Some individuals may also lack phenotypic data, as it is in the case when the problem is predicting yet-to be-observed outcomes.

The Hy-BLUP setting is more general than that considered in ssGBLUP. In the latter, all individuals must be members of a known genealogy and only a fraction has molecular data. ssGBLUP exploits the flexibility conferred by mixed linear methodology and has received much attention in animal breeding. Essentially, breeding values of non-genotyped individuals are imputed from those that have markers using an approximation to a conditional distribution. Legarra et al. (2009), Aguilar et al. (2010) and Christensen and Lund (2010) present central ideas of ssGBLUP and Fernando et al. (2014) describe a Bayesian treatment. Other studies include ad-hoc methods or adjustments to introduce “compatibility” of for circumventing singularity of some intervening matrices, e.g., Christensen et al. (2012) and López-Cruz et al. (2015). These adjustments are tweaks of the central prediction machine that flows directly from the distribution theory used by Aguilar et al. (2010).

Hy-BLUP shares the spirit of ssGBLUP but it also accommodates individuals lacking pedigree that may have been genotyped for forensic, commercial or traceability reasons. For instance, a meat packing plant takes DNA samples from animals without known lines of descent. Hy-BLUP uses a direct Bayesian logic and does not involve approximations, contrary to ssGBLUP. The basic notion is one of combining independent prior opinions whose degree of precision is conveyed by the inverse of covariance matrices of Gaussian distributions. Further, additional weights may (may not) be placed on the sources of similarity (pedigree and molecular) among individuals, providing an extra tool for tuning predictive performance.

### Motivation for Hy-BLUP

Robertson (1955) cast the problem of estimation of breeding values as follows. Say it is known that the distribution of breeding values in a population is 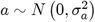 where 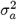 is the additive genetic variance. The distribution represents a statement of uncertainty about the breeding value of an individual, prior to collecting any phenotypic information. An individual *i* is drawn at random from the population and measured *n*_*i*_ times, with the model of its record *j* being *y*_*ij*_ = *μ* + *a*_*i*_ + *e*_*ij*_, where *μ* is a location parameter and 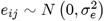 is a random residual. Suppose 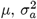 and 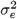 are known, so one can work with the deviations *y*_*ij*_ − *μ*.

Robertson stated that there are two independent sources of information about *a*_*i*_. One derives from knowledge of the population: the expected breeding value is 0 and the precision is 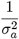. The second source is the least-squares estimator (maximum likelihood under normality) of *a*_*i*_: 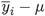, having precision 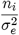. The total precision after (posterior to) observation is 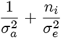. The two statements 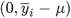 can be combined using their respective precisions as weights into the weighted average

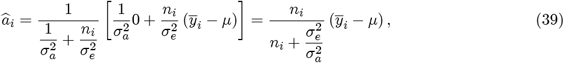

which yielding a textbook formula for “estimated breeding value” in this simple setting. Here, the “regression” 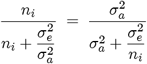, is sometimes called “heritability of a mean”, and 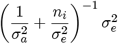 is the “prediction error variance” or variance of the conditional distribution 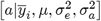, which is the posterior variance in a Bayesian context. Dempfle (1977) showed how this reasoning could be used for deriving BLUP. In the simple setting of Robertson (1955), 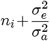 plays the role of the “coefficient matrix” in the mixed model equations (proportional to posterior precision), and 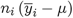 is the counterpart of the vector of right-hand sides. Robertson’s solution is a special case of equations (5) and (6).

The Bayesian approach enables combination of various prior opinions. Instead of using a sole source of prior information, a joint prior “opinion” derived from several sources can be arrived at by taking each opinion as a single data point. An example is in Press (1989) where expert opinions were combined and used to construct a prior probability distribution of a nuclear war. In a less ominous context, a Bayesian may have a prior opinion about a breeding value derived from an infinitesimal model while other Bayesian may derive the prior from a genome-based model. Clearly, these two priors differ in variance: the unknown genetic value is the target random variable but one opinion uses 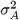 as prior variance while other takes 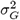. These two Bayesians are willing to combine their opinions but encounter the difficulty of eliciting a joint prior distribution of unobservable infinitesimal and genomic values. Then, they agree to treat their opinions as independent, perhaps overstating the strength of prior knowledge. They add up the precisions provided by the pedigree (reciprocal of the pedigree-captured variance) and by the genomic (reciprocal of the genomic variance) data. Estimates of these two variances are now abundant for many species of plants and animals that have been genotyped during the last two decades (Gondro et al. 2013; Isik et al. 2017; Ahmadi and Bartholome 2022), so the precision conveyed by a joint opinion can be derived readily. The idea is similar to the concept of information in likelihood. For instance, suppose observations 1 and 2 have a common mean but variances *V*_1_ and *V*_2_. Fisher’s information about the mean from the sample is 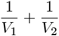; the variance is 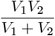, which becomes 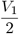 if the variance is homogeneous; here, 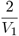 is the information (likelihood) or precision (Bayesian) under the homogeneity assumption.

Returning to Robertson (1955), write the prior precision about a breeding value *u* as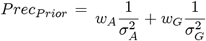, where *w*_*A*_ and *w*_*G*_ are arbitrary weights. A posteriori, 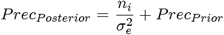 and the estimated breeding value is now

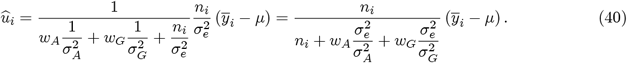

The linear model implicit in this treatment of Robertson’s view is 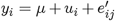, where *u*_*i*_ is a random variable with variance

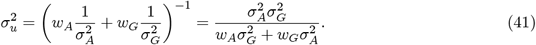

Normatively, *w*_*A*_ = *w*_*G*_ = 1, in which case 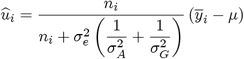 and 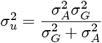.

*Hy*−*BLUP* flows from a matrix generalization of the concept. As given in (26) the variance-covariance matrix of breeding values in *Hy* − *BLUP* is

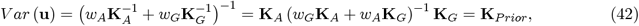

which is the matrix counterpart of (41). For simplicity, let *β* =0 and suppose 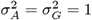 so reproducing kernel Hilbert spaces regression representations of ABLUP, GBLUP, ssGBLUP of and *Hy* − *BLUP* (Gianola and van Kaam 2008; de los Campos et al. 2009) are

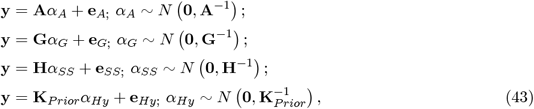

where the *α*′ *s* are regression coefficients of phenotypes on kernel matrices (e.g., **H**) and the **e**′*s* are residuals. Both ABLUP and GBLUP are linear regressions of phenotypes on pedigree and genomic relationship matrices, respectively. However, ssGBLUP and Hy-BLUP are not linear in **A** or **G** as seen from the forms of **H** in (23) and **K**_*Prior*_. Hence **H** and **K** involve nonlinear features on similarities, a property that perhaps confers these methods the ability of capturing more forms of complex gene actions, even though **A** and **G** encode additive similarities.

### Conceptual differences between Hy-BLUP and ssGBLUP

The innovation in HY-BLUP relative to ssGBLUP is in the prior adopted. The assumptions 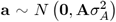 and 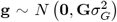 can be viewed as two different statements of prior uncertainty about an unknown additive value **u**, defined at the level of unknown QTL and with variance 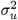. In Hy-BLUP, the two pieces of prior information are combined (considering the various patterns in which the data appear) as if they were independent. This assumption may lead to an overstatement in prior precision (smaller variance and more shrinkage) because *a* and *g* must have a positive covariance. Since these two variables try to capture the same “causal variable”, *u*, they must have a joint distribution albeit difficult to specify. Hy-BLUP has the flexibility of accommodating complex data structures and the prior should be gradually over-ridden as information accrues. The computations required for forming the Hy-BLUP mixed model equations are similar to those in ssGBLUP but the logic of the latter method is different.

From statistical theory (we use scalars for simplicity), the joint prior distribution of two pedigree breeding values with *E* (*a*_1_)= *E* (*a*_2_)= 0, homoscedasticity and normality assumptions has density

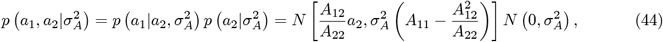

where *A*_11_, *A*_12_ and *A*_22_ are elements of the 2 × 2 “numerator”relationship matrix. The model above implies that 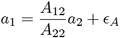, where 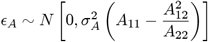 is a residual. Likewise, the density of the joint distribution of the genomic breeding values *g*_1_, *g*_2_ of the two individuals is

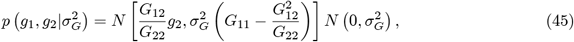

where *G*_11_, *G*_12_ and *G*_22_ are elements of a 2 × 2 genomic relationship matrix, so 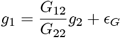, where 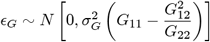. In ssGBLUP (e.g., Aguilar et al. 2010), the assumption made is that the two breeding values have joint density

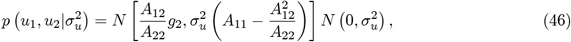

which implies that 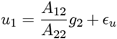 with 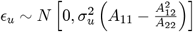 taken as distributed independently of *g*_2_. Note that 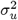 is neither 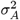 nor 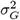. Save for the variance parameter (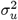 replaces 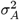), *ϵ*_*u*_ is as *ϵ*_*A*_ in the pedigree-based model. This structure and independence assumption lead to the marginal variance

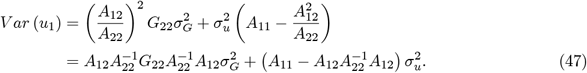

If one takes 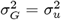, then 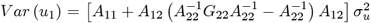, as given in matrix form in (23). These assumptions made in ssGBLUP are not made in Hy-BLUP.

How can missing pedigree information be accommodated in ssGBLUP? Take the situation in (12), where individuals with breeding values **u**_2−_ have markers but cannot be traced to relatives in the population. The imputation used in ssGBLUP would be

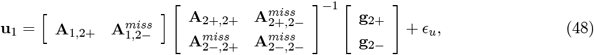

where “*miss*” denotes “missing. Clearly, there are parts that cannot be filled. Similarly, the **H**^−1^matrix required for forming the mixed model equations in (4) would have the (symbolic) form

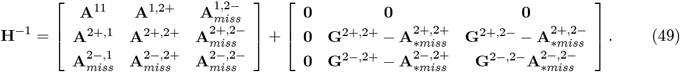

Here, the **G** sub-matrices with superscripts are partitions of the inverse of the genomic relationship matrix of individuals with markers, and the **A**_*_ sub-matrices are portions of the inverse of the numerator relationship matrices. Since there are missing parts, **H**^−1^ cannot be formed properly.

A possible solution could make use of the “memory property” of Bayes theorem. Ignoring temporarily the pedigree information, the joint posterior distribution of the genomic will have mean vector (GBLUP)

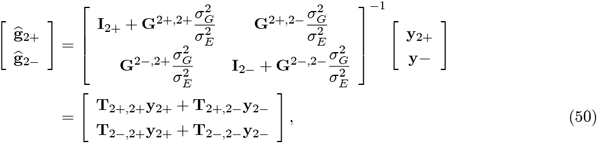

where

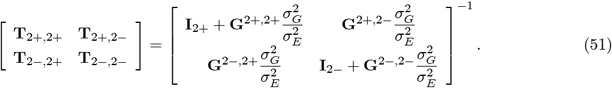

Hence, the posterior (to phenotypes of individuals with marker data) distribution of the genomic breeding values in set 2+ is 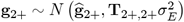. Next, use this information in ssGBLUP and form the mixed model equations as

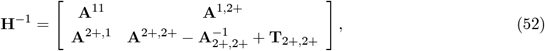

and the right hand-sides (assuming no fixed effects in the model) would be

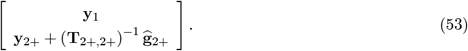

Hence, ssGBLUP can be adapted to accommodate individuals having markers but with missing pedigree information. HY-BLUP seems simpler, at least in this respect,

### Tuning weights

In our evaluation of *Hy* −*BLUP*, a grid of fixed values of *w*_*A*_ was chosen and examined in terms of out-of-sample predictive performance, given the variance components. Can these weights be estimated somehow? Consider a random effects Hy-BLUP setting where the marginal distribution of the data leads to the likelihood function

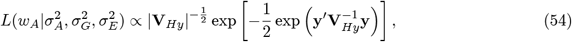

with *w*_*A*_ =1 − *w*_*G*_ and *w*_*A*_ + *w*_*G*_ = 1. Here

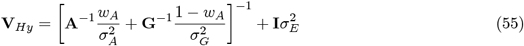

where **A**^−1^ and **G**^−1^are formed as in (9) and (10), respectively. There is an infinite number of combinations of values of *w*_*A*_ and 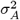 that give the same ratio, and similarly for 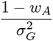 there is an identification problem and the parameters *w*_*A*_, 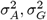 are not individually estimable. Given *w*_*A*_, however, the variance components can be inferred from the likelihood, as discussed later.

If there are reliable estimates of 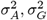 and 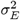, an analytical Bayesian approach can be used to assess the relative plausibility of a discrete set of values of *w*_*A*_ over a grid of *m* values. Let 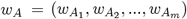 with each element assigned equal prior probability 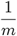. The (conditional) posterior distribution of *w*_*A*_ is

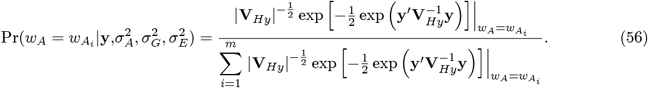

To illustrate, conditional posterior distributions of the weight placed on pedigree (*w*_*A*_) were computed with (56) at four combinations of values of the variance parameters 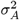 and 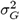 with 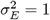, using **A** and **G** from the entire data set. Values of *w*_*A*_ ranged from 0 to 1, at increments of 0.001; the prior probability of each value was therefore 10^−3^. The posterior distributions are displayed in Figure 10. The posterior distributions differed from the prior distribution, which would be a horizontal line placed at 10^−3^ on the Y-axis. The posterior distributions tended to assign larger probabilities to values of *w*_*A*_ lower than 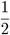 and distributions got less sharp as the ratio 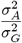 increased; in the most extreme case with 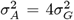 (blue curve), values of *w*_*A*_ larger than 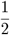 were assigned more posterior probability but the curve was quite flat. Hence, given the variance parameters, some Bayesian learning about the weight on pedigree takes place. Note, however, that the values of 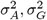 (acting as hyper-parameters here) were influential, meaning that learning about *w*_*A*_ was weak.

**FIGURE 10.**
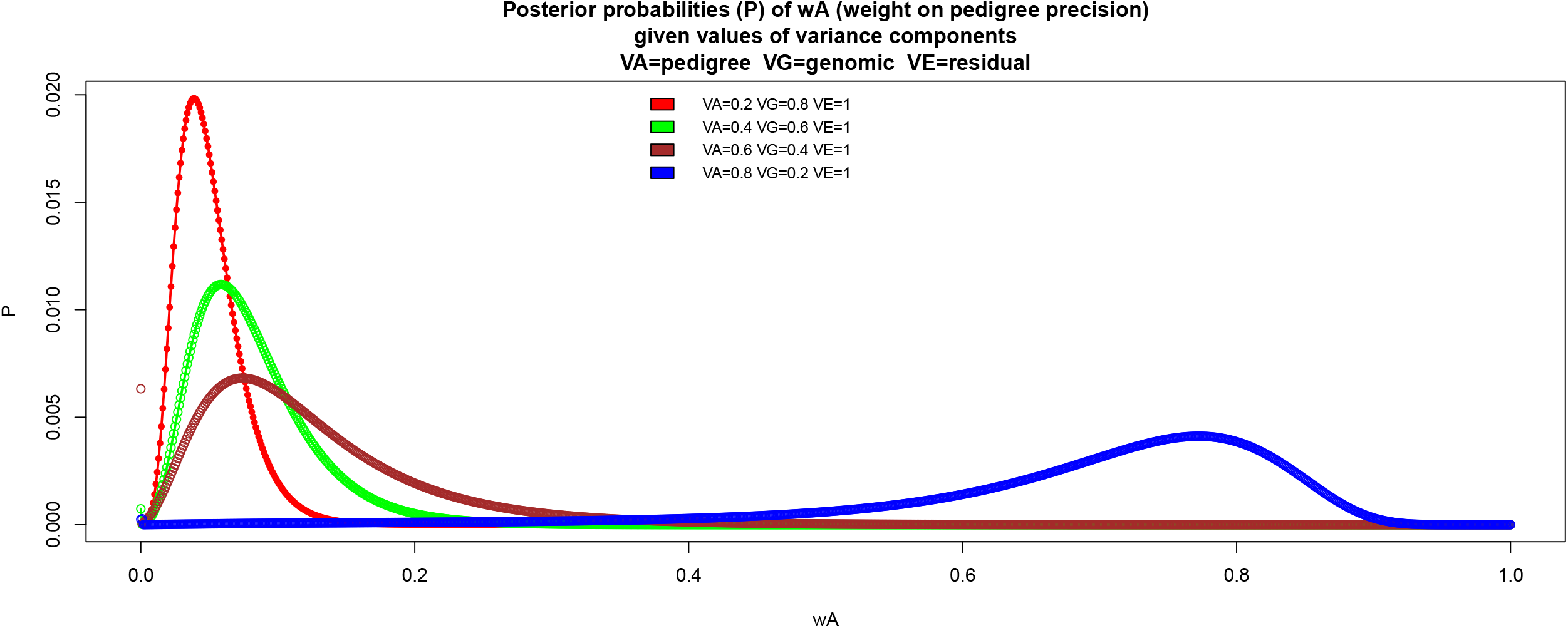
Posterior probabilities of weight placed on value conditionally in variance component values. VA=variance captured by pedigree. VG= variance captured by molecular similarity. VE= residual variance.

While learning *w*_*A*_ may have little inferential added value because of the “confounding” with 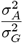 the weights placed on precision from pedigree or markers may impact predictive ability differentially. An “estimate” of weight derived from training data may not deliver the best predictive performance (e.g., Gianola 2021; Reinoso-Peláez et al. 2022). For this reason, we favored calibrating the weights in terms of predictive ability.

### Learning variance parameters in “normative” Hy-BLUP

A distinctive aspect of Hy-BLUP is that two variance components 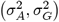 govern the prior distribution of breeding values. Can these two parameters and 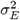 be estimated from the likelihood function? This question is examined next from the perspective of the EM algorithm, where E stands for “Expectation” and M for “maximization” (Dempster et al. 1977; Little and Rubin 1987; Sorensen and Gianola 2002). We take *w*_*A*_ = *w*_*G*_ =1 because *w*_*A*_ (*w*_*G*_) and 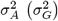 are not identified in the likelihood, e.g., there is an infinite number of pairs of *w*_*A*_ and 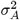 that produce the same ratio 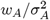.

The “complete data” log-likelihood for a Hy-BLUP(1,1) random effects model is

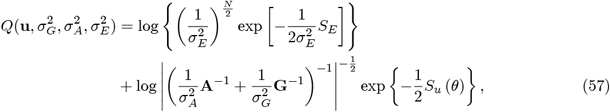

where *S* = (**y** − **Zu**)′ (**y** − **Zu**) and 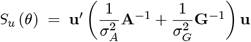 are quadratic forms on the breeding values **u**, and 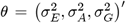 is the vector of variance parameters. Note that *S*_*E*_ does not depend on *θ* while *S*_*u*_ (*θ*) involves the genetic variances. Starting at some initial point, let the variance components at iteration *t* take values 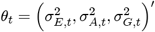. Define

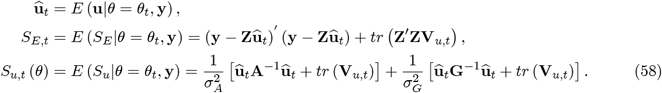

Above **û**_*t*_ and **V**_*u,t*_ are calculated as

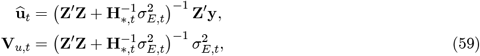

where 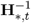 is as in (16), that is

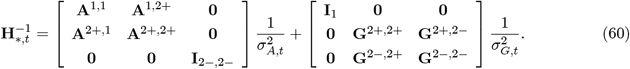

Using the preceding expressions, the *Q* function at iteration *t* is estimated in the E-step of the algorithm as

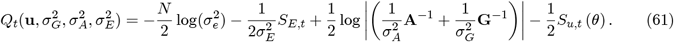

The M-step consists of taking first derivatives of 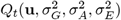 with respect to the unknown variance parameters, setting to zero. and solving. Here

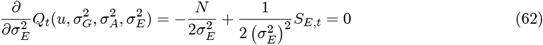

produces as solution for the next iterate

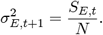

The derivatives with respect to genetic components are more involved because of the determinantal expressions. Here

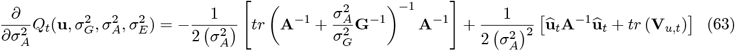

and

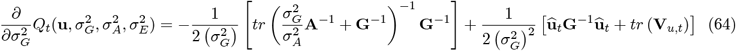

Setting the “genetic” derivatives to zero does not produce an explicit system and rearrangement produces

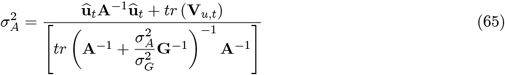

and

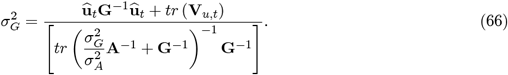

An ad-hoc way of continuing the iteration consists of replacing 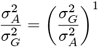 by the value of the ratio at round *t*, yielding new values 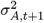 and 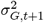

In a nutshell: the Hy-BLUP setting allows disentanglement of pedigree and genomic sources of variation. Future work should investigate the topic in more detail, including algorithms based on second derivatives or Bayesian methods.

### Multiple-trait extension

Suppose there is a set of commercial animals slaughtered at a packing plant; such animals have been measured for a carcass traits and some have DNA samples, but cannot be traced to relatives in the population. Two questions of interest might be: 1) what is the genetic lag between commercial animals and those in the breeding nucleus for traits that are routinely recorded in breeding schemes (e.g., weaning or yearling weights in beef cattle)? 2) Can the DNA information be used to “propagate” carcass information (from destroyed animals) to those in the breeding population? Hy-BLUP provides an answer to both questions.

Let **y** and **z** denote vectors of measurements for a production trait and for a carcass measure, e.g., yearling body weight and “cold” carcass weight, respectively. Animals with records for the production trait may have a complex pattern of pedigree and genomic information; the animals with carcass traits will be assumed to posses markers but no pedigree. The breeding values for all animals, using the notation in (7) are

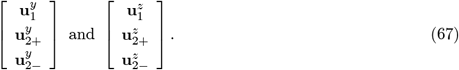

Recall that animals in set (2−) have markers but are disconnected from the population, pedigree-wise. Let the 2 × 2 pedigree-based and genome-based covariance matrices between traits be

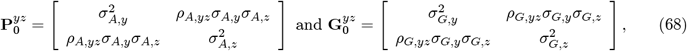

respectively. Here 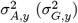 are the pedigree and genomic variances for “trait *y*” and likewise for “trait *z*”; the *σ*′*s* are genetic standard deviations. The genetic correlations between traits from pedigree and markers are *ρ*_*A,yz*_ and *ρ*_*G,yz*_, respectively. A simple extension to the bivariate situation of developments leading to (12) is

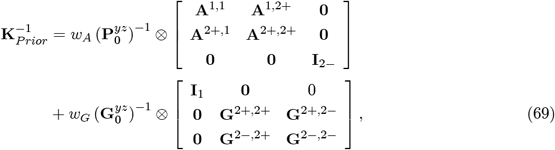

where *w*_*A*_ = *w*_*G*_ = 1 if the normative version is adopted. Since *y* and *z* are observed in different animals, the residual correlation can be assumed null, that is, the residual covariance matrix is 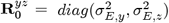. Hy-BLUP predictions of breeding values of all animals for each of the two traits, as in (67), can be obtained by solving standard bivariate mixed model equations, with 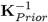 taken as covariance matrix of breeding values. Thus, all animals in the breeding population would have predictions of their genetic worth for the carcass trait indirectly from measurements obtained in packing plants via genomic similarities conveyed by DNA, assuming parameter estimates are available, especially the genetic correlations (pedigree and genomic) between the two traits.

A special case is when slaughtered animals, while having DNA samples, have not been recorded individually for the production trait. Here, via” genomic connectedness”, one can obtain Hy-BLUP of their breeding value for the production trait. These predictions are not useful for selection decisions and would be imprecise, but lead to an estimate of “genetic lag” between nucleus and commercial animals. Using Hy-BLUP, calculate the average estimate of breeding values for a given cohort of animals and compare with the mean estimated breeding value in the commercial animals. By dividing the difference between the averages by an estimate of genetic trend, an approximation to the genetic lag in time units can be obtained.

## 6 Conclusion

Relative to SS methodology, the novelty of HY-BLUP is that predicted breeding values can be obtained for individuals without pedigree, with or without records, that have been genotyped. The HY-BLUP procedure retains the logic and flexibility of BLUP but it is the outcome of Bayesian reasoning.

The behavior of Hy-BLUP was examined using a wheat data set and the method was compared with ssGBLUP. At least under the conditions of our study, differences between ssGBLUP and HY-BLUP, even though detectable, were small. There were situations, however, in which Hy-BLUP performed slightly better than ssGBLUP. A possible reason for this difference in favor of Hy-BLUP is because it serves as a proxy for a multi-kernel procedure, a form of model averaging that can produce a better mean-squared error performance (Sorensen and Gianola 2002; de los Campos et al. 2010). Another reason is that, when interpreted as kernel regression, Hy-BLUP makes non-linear features out of “linear similarities”, which may confer flexibility for capturing more complex genotype-phenotype maps. Additional studies of Hy-BLUP with other data sets seem warranted and methods for (co) variance structure estimation developed accordingly.

In the discussion, we gave a scalar motivation of the procedure and showed how the notion extends to the multi-trait situation. Given the variance components, weights can be learned from the data, although to a limited extent. However, in the “normative” Hy-BLUP (*w*_*A*_ = *w*_*G*_ = 1) weights are not necessary and this specification may have a better performance from the perspective of prediction, the main focus of our paper. We discussed conceptual differences with ssGBLUP and showed that the genomic and pedigree-based variances employed in Hy-BLUP can be estimated from the data at hand.

Employing a wheat grain yield data set, an empirical comparison between the two methods did not produce conclusive evidence of practically meaningful differences between the two procedures. Additional studies with other species and traits may be able to detect differences, a finding that would be consistent with the view that there is no such thing as a universally best predictor (Gianola 2021).

## 7 Acknowledgments

Gustavo de los Campos (Michigan State University) provided the *R* code for maximum likelihood estimation of the variance components.

